# Fungal Mineral Weathering Mechanisms Revealed Through Direct Molecular Visualization

**DOI:** 10.1101/2021.10.01.462718

**Authors:** Arunima Bhattacharjee, Odeta Qafoku, Jocelyn A. Richardson, Lindsey N. Anderson, Kaitlyn Schwarz, Lisa M. Bramer, Gerard X. Lomas, Daniel J. Orton, Zihua Zhu, Mark H. Engelhard, Mark E. Bowden, William C. Nelson, Ari Jumpponen, Janet K. Jansson, Kirsten S. Hofmockel, Christopher R. Anderton

## Abstract

Soil fungi facilitate the translocation of inorganic nutrients from soil minerals to other microorganisms and plants. This ability is particularly advantageous in impoverished soils, because fungal mycelial networks can bridge otherwise spatially disconnected and inaccessible nutrient hotspots. However, the molecular mechanisms underlying fungal mineral weathering and transport through soil remains poorly understood. Here, we addressed this knowledge gap by directly visualizing nutrient acquisition and transport through fungal hyphae in a mineral doped soil micromodel using a multimodal imaging approach. We observed that *Fusarium sp. DS 682*, a representative of common saprotrophic soil fungi, exhibited a mechanosensory response (thigmotropism) around obstacles and through pore spaces (∼12 µm) in the presence of minerals. The fungus incorporated and translocated potassium (K) from K-rich mineral interfaces, as evidenced by visualization of mineral derived nutrient transport and unique K chemical moieties following fungal induced mineral weathering. Specific membrane transport proteins were expressed in the presence of minerals, including those involved in oxidative phosphorylation pathways and transmembrane transport of small molecular weight organic acids. This study establishes the significance of fungal biology and nutrient translocation mechanisms in maintaining fungal growth under water and nutrient limitations in a soil-like microenvironment.

## Introduction

Biotic mineral weathering is critical in impoverished soils where nutrients can be bound to minerals and unavailable to plants and other organisms. Some soil fungi are adept at extraction of micronutrients from soil minerals by weathering rock material to mineralize elemental potassium (K), iron (Fe), manganese (Mn), calcium (Ca), and other inorganic nutrients^1–3^. These fungi release elemental nutrients from rock derived minerals either (i) indirectly, by secreting low molecular weight organic acids into the soil microenvironment, or (ii) directly, by exerting physical forces at the hyphae-mineral interface^4^. Some mycorrhizal fungi then transfer the mineral derived nutrients to host plants in exchange for carbon^5–8^. Several instances of both ectomycorrhizal and arbuscular mycorrhizal fungal weathering of P and K containing soil minerals have been documented, where the main mechanism of weathering is by dissolution with organic acids^9–12^. Specifically, mycorrhizal associations with plants improve K uptake through fungal induced mineral weathering and transport of K to their host plants^13–15^. Saprotrophic fungi can also contribute to soil mineral weathering^16–20^, although less is known about the underlying mechanisms when compared to mycorrhizae. A study with the saprotrophic fungus *Aspergillus niger*, found evidence of direct and indirect mineral weathering in liquid cultures containing mineral grains^21^.

The above mentioned studies have demonstrated that widespread fungal hyphal networks on different soil mineral surfaces facilitate weathering of minerals and transport of mineral-derived nutrients through the complex soil landscape^22–24^. However, fungal induced mineral weathering and nutrient transport is a spatially heterogeneous process in nature, and bulk soil measurements do not provide a fully representative understanding of fungal hyphal interactions at the mineral surface within soil systems. Moreover, the filamentous and microscopic nature of fungal hyphae requires investigation of mineral weathering at the hyphae-mineral interface, and the ability to resolve weathering processes spatially at these microbial scales could provide more comprehensive insights into nutrient transport at the ecosystem scale. Further complicating the matter, diffusion of fungal exuded organic acids in saturated soils can dramatically alter the heterogeneity of mineral weathering processes^25^. As such, the spatial heterogeneity of mineral weathering processes in soil presents unique challenges in experimentally identifying specific fungal interactions within the complex soil habitat. Consequently, mineral weathering and transport mechanisms in fungi have not been spatially analyzed, in part due to the lack of analysis platforms that allow spatial visualization and analysis of fungal induced mineral weathering. At the bulk soil scale, fungal induced mineral weathering has been extensively studied in mycorrhizal fungi^9–11^. However, mechanistic investigations of the contributions of mycorrhizal fungal to mineral weathering are difficult due to their strong associations with host plants for survival. Investigating mineral weathering in saprotrophic fungi could demonstrate mineral derived nutrient transport mechanisms unique to fungal communities in the absence of a plant host.

Here, we developed a novel mineral doped soil micromodel platform that provided unprecedented spatial visualization into the molecular mechanisms underpinning thigmotropism, mineral weathering, and nutrient transport by a soil saprotrophic fungus, *Fusarium sp. DS 682*^26^. Using this platform, we demonstrate that this species can degrade and acquire minerals through extraction of K by an indirect mineral weathering mechanism in an environment that mimics select soil physical and chemical properties. The results of this study provide a microscale level understanding of fungal weathering of soil minerals, K^+^ and Na^+^ transport, and K speciation in minerals after fungal growth.

## Results

### Development of the mineral doped soil micromodel platform for probing fungal induced mineral weathering

We developed a mineral doped soil micromodel (Fig. 1a) that simulates a weathered soil environment, where key nutrients (e.g., K, Fe, Ca, Mn) were not readily bioavailable because of their strong associations to minerals. The soil micromodel features an evenly spaced matrix of pillars that mimics pore spacing found in the soil environment^27^, and the device was fabricated with polydimethylsiloxane (PDMS) doped with minerals. To build the mineral doped soil micromodel, we used a natural kaolinite powder that contained 7% K-feldspar and 4% mica— phase components that are common in minerals present within soil^28^. Elemental analysis of the natural kaolinite showed an abundance of essential nutrients such as K, Na, Mg, and Ca (Fig. 1b), in addition to Al and Si. These essential nutrients originated from the K-feldspar and mica in the kaolinite (Fig. S1). The use of a natural mineral enabled interrogation of the release and transport of mineral nutrients by the fungal hyphae under controlled environmental conditions. The minerals embedded in the surface of the microchannel in the soil micromodel also provided a direct contact interface between the fungus and embedded minerals (Fig. 1c and S2).

**Fig. 1.**
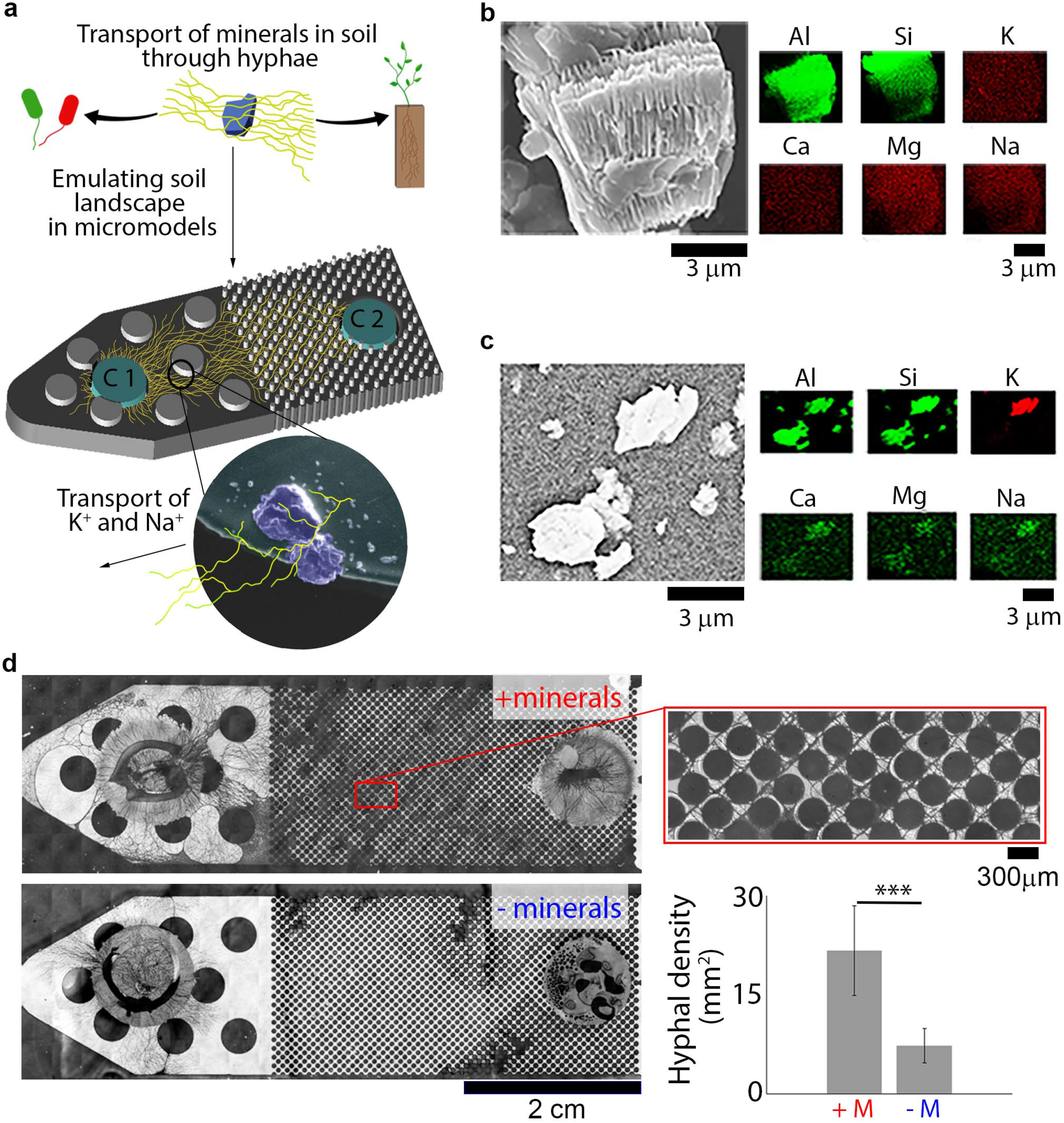
Fungal growth during water and nutrient limitation is contingent on the presence of soil minerals with accessible nutrients. **a)** A schematic of the soil micromodel used for this study, which emulates natural soil pore spacing and mineralogy. Fungal growth in a water and nutrient limited environment are enhanced in mineral doped micromodels due to extraction of K^+^ and Na^+^ from minerals through mycelia. The PDA plugs, ‘C1’ and ‘C2’, are the only nutrient sources present in the micromodel, and the fungus was inoculated on C1. **b)** Energy dispersive X-ray (EDX) images of the powdered minerals used to dope micromodels with minerals shows the presence of essential elements, such as K, Ca, Mg, and Na, which could be used by the fungus for growth in this water and nutrient limited microenvironment. **c**) EDX images of the same mineral powder embedded in the PDMS matrix, which demonstrates their accessibility in these micromodel systems. **d)** PDMS mineral doped micromodel with minerals (top, + minerals) shows enhanced fungal growth, whereas fungal growth is limited in PDMS micromodels without minerals (bottom, - minerals). We observed thigmotropic response in the mineral doped micromodel shown in the zoomed-in image on right within the red square. Quantitative fungal growth comparisons in nutrient limited soil micromodels with (+M) and without (-M) minerals shows a significant increase in hyphal density in the presence of minerals (bar graph, bottom right). *** Denotes a p value < 0.0001, image analysis method is described in Fig. S5.

The soil micromodel was inoculated with *Fusarium sp. DS 682*, a saprotrophic fungus that is common in grassland soils^26^. The fungus was inoculated at one end with a potato dextrose agar (PDA) plug (Fig. 1a, location C1), while a second axenic PDA patch (Fig. 1a, location C2) was provided at the other end of the device. The mineral grains embedded in the micromodel were the only source of nutrient, such as K, Na, Ca (Fig. 1a) between the PDA plugs (C1 and C2) at either end of the device, creating a nutrient impoverished condition. Fungal driven weathering of the minerals was thus required for making the mineral nutrients bioavailable. A water limitation stress was simulated in the soil micromodel by creating an unsaturated channel to keep the PDA sources apart, preventing diffusion of nutrients between PDA and mineral nutrient (Fig. 1a) through liquid media in the microchannel. Therefore, fungal contact with minerals was required to induced weathering of mineral surfaces for extraction of nutrients. These stresses generated an environment for studying fungal degradation and transport of mineral derived nutrients.

### Mineral availability regulates fungal growth in nutrient limited environments

The micromodel system provided an unprecedented view of fungal mineral weathering mechanisms using several spatially resolved characterization techniques, such as optical microscopy, electron microscopy, secondary ion mass spectrometry, and X-ray spectromicroscopy. The PDMS-glass configuration of the micromodels was reversibly bonded such that the PDMS mineral micromodel side could be separated from the glass coverslip for characterizing fungal mineral weathering using advanced imaging techniques.

We used optical microscopy to image the movement of the fungal hyphae from the inoculation site (C1) through the porous environment (12-150 µm pore spaces) to access the axenic nutrient (PDA) pool (Fig. 1a, C2). We observed that fungal mycelia grew extensively in the presence of minerals and exhibited thigmotrophic movement around micro- and macro-pore spaces that mimicked natural soil pores (Fig. 1d). On the contrary, the absence of soil minerals in the microenvironment significantly inhibited fungal growth (Fig. 1d), where fungi grew only around the nutrient plug at the inoculation site (C1). Further tests suggest that this increase in hyphal density in microporous environments mimicking soil were specifically regulated by the fungal-mineral interactions. Notably, the porosity of the micromodels without minerals did not significantly alter hyphal growth, and fungal hyphal biomass was observed in micromodels with larger pore spaces (600 µm). However, growth ceased when the pore size decreased to 5 µm (Fig. S4a), demonstrating that thigmotrophic movement was at least partly dependent on pore size. Additionally, thigmotrophic movement could not be attributed to the surface chemistry of the PDMS, as micromodels with either hydrophilic or hydrophobic surfaces resulted in fungal growth comparable to micromodels without minerals (Fig. S4b). Combined, these results illustrate the role of minerals in fungal hyphal thigmotrophic movement within microporous environments mimicking soil.

### Direct visualization of fungal nutrient uptake from minerals and K speciation by fungal hyphae

We observed an increase in the relative concentrations of K^+^ and Na^+^ in fungal hyphae grown in mineral doped soil micromodels compared to mineral free micromodel controls by time-of-flight secondary ion mass spectrometry (ToF-SIMS) imaging (Fig. 2a). The micromodels with and without minerals had the same amount of PDA, which contained minimal amounts of K and Na. Therefore, any increase in relative concentrations of K^+^ and Na^+^ in the fungal hyphae in the mineral doped micromodels was attributed to uptake from the K-feldspar and mica minerals. We did not observe an increase in other mineral nutrients detected within mineral grains shown in Fig. 1b and c (e.g., Ca, Mg) into the fungal hyphae.

**Fig. 2.**
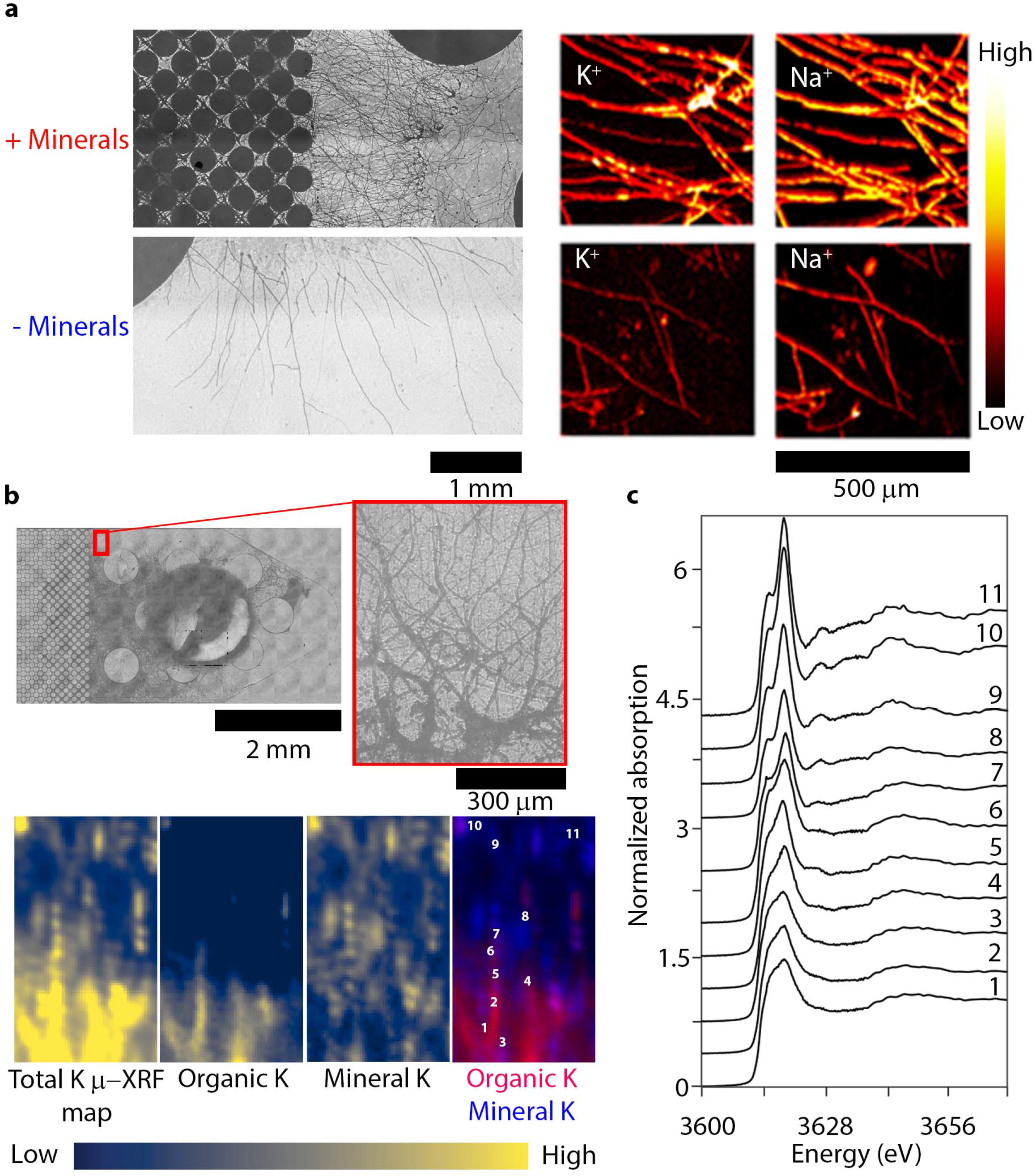
Direct evidence of fungal hyphal uptake and organic complexation of essential nutrients from degrading minerals in soil micromodels. **a)** Optical microscopy images of fungal growth (left) and time-of-flight secondary ion mass spectrometry (ToF-SIMS) analysis (right) of fungal hyphae grown in micromodels with and without minerals. ToF-SIMS results demonstrate an increased uptake of K^+^ and Na^+^ ions along fungal hyphae in micromodels containing minerals in comparison to those grown in a mineral-less environment. **b)** Optical image of mineral doped micromodel showing the µ-XRF mapping region in red box with corresponding µ-XRF maps. Maps of (i) total K (above the K K-edge at 3627 eV), (ii) organic complexed K, (iii) mineral bound K, and (iv) a dual color plot of organic complexed K (red) and mineral bound K (blue) with numbers corresponding to locations of XANES spectra in **c**. Maps of organic K and mineral K were created from fitting of multi-energy maps with end-member spectra. The hyphae rich region in the optical image corresponds to organic complexed K in the XRF maps and shows different forms of K exists in the micromodel, as a result of fungal growth **c)** XANES spectra from the locations numbered in XRF maps in **b**. Spectra show a change from organic K (numbers 1-6) dominated to mineral bound K (numbers 7-11) across the mapped region. In spectra representing mineral bound K (7-11), there are shifts in the peak energy, demonstrating changes in K mineral chemistries precipitated from fungal degradation of minerals in the micromodel surface. The LCF of the spectra 1-11 showing the % of mineral and organic complexed K is shown in Fig. S8.

We hypothesized that enhanced fungal growth in the presence of minerals was due to uptake of nutrients by hyphae from minerals in the micromodel through mineral weathering by the release of fungal organic acids. Consequently, we expected to observe differences in K chemistry in mineral interfaces and K^+^ transported in fungal hyphae from K minerals. Using multi-energy micro-X-ray fluorescence imaging (µ-XRF) around the potassium K-edge, combined with potassium X-ray Absorption Near Edge Structure (XANES) spectroscopy we found evidence of K^+^ transport from minerals through the fungal hyphae (Fig. 2b and c). Specifically, we identified two distinct K chemistries in the mineral doped soil micromodel as a result of fungal growth: (i) inorganic mineral-bound K (referred to as ‘*inorganic K*’), and (ii) unidentified organic, possibly hyphal adsorbed, K (referred to ‘*organic K*’) located along distinct paths (Fig. 2b). The distinct paths of the *organic K* coincided with elevated S and P abundances that are indicative of fungal hyphal biomass (Fig. S6). The *organic K* was observed at the fungal inoculation point and in fungal hyphae grown with and without minerals, suggesting that this form of K is highly present in fungal hyphae.

The extensive fungal induced K speciation of the minerals was probed using spatially defined XANES spectra within the soil micromodel (Fig. 2c). Here, mineral K showed the distinct pre- and post-edge features (Fig. 2c, spectra 7-11), which were absent in the *organic K* spectra collected from fungal biomass growing in the same region (Fig. 2c, spectra 1-6). Interestingly, we observed peak shifts in energy of the spectral features within inorganic K spectra (Fig. 2c, 7-11), while the peaks for *organic K* spectra (Fig. 2c, 1-6) did not exhibit energy shifts. Mineral weathering by different fungal derived organic acids resulting in distinctive mineral K bonding environments could contribute to these shifts in energy observed for the pre- and post-edge features in inorganic K spectra (Fig. 2c and S8). In addition, a linear combination fitting of end-member organic K and mineral XANES spectra (Fig. S7) to the experimental micromodel K XANES spectra indicates discrepancies in the fits of spectra 7-11, which alludes to a change in the mineral surface chemistry (Fig. S8).

Furthermore, hyphal trails that specifically tracked with the location of mineral grains rich in K were revealed by imaging the micromodel surface after fungal biomass was removed (Fig. S9). This imaging was made possible because areas where the hyphae attached to the surface of the micromodel were enriched in carbon and directly observable by scanning electron microscopy (SEM) imaging. These images suggest the potential that fungi can sense and extract mineral derived nutrients based on their requirement of a specific nutrient, K in this case. We did not detect formation of previously reported micro-tunnels, cavities, or micropores on the mineral grain surfaces, as observed during direct mineral weathering by ectomycorrhizal fungi^4,29^. Together, our results showing transport of K and Na through hyphae, K speciation in micromodel minerals after fungal growth, and the absence of micro-tunnels on micromodel mineral grains suggest that the mineral derived nutrient extraction by *Fusarium* hyphae is via an indirect weathering mechanism.

### P-type ATPases and transmembrane transporter proteins enable K transport from minerals into fungal biomass

Indirect mineral weathering occurs through fungal secretion of different organic acids that are used to dissociate and uptake mineral-bound nutrients^4,30,31^. As such, we hypothesized that *Fusarium sp. DS 682* employed specific fungal transporter proteins that facilitate passage of organic acids to increase assimilation of mineral derived nutrients into the nutrient limited micromodel environment. Accordingly, we expected to see such transporters expressed in greater abundance in the presence of minerals. To test this hypothesis, *Fusarium* sp. *DS 682* was grown in direct contact with a thin layer of natural kaolinite minerals on the surface of PDA to collect biomass for proteomics. The fungal growth was much faster in plates with minerals than without minerals (Fig. S10). Proteins were extracted and identified by mass spectrometry, resulting in a total of 4313 fungal proteins detected (Fig. 3a and S11), where several families of transporters were expressed under both conditions, with minerals (+M) and without minerals (-M). These proteins included the major facilitator superfamily (MFS) transporters, the solute carrier (SLC) transporter, ATP-binding cassette (ABC) type transporters, and Aquaporins (Fig. 3b). However, specific transporter proteins were enriched under both conditions. We observed 34 transport proteins significantly enriched under -M condition, while 15 transport proteins were significantly enriched under +M condition. Among these transport proteins, we also observed several membrane ATPases that are essential for transmembrane transport of nutrients expressed under both conditions. Specifically, several key subunits of the oxidative phosphorylation pathway, such as vacuolar membrane V-type ATPases and mitochondrial membrane F-type ATPases were detected under both conditions (Figs. 3c and S12). Further, we observed a P-type ATPase in the oxidative phosphorylation pathway for transmembrane transport enriched in the +M condition alone (Figs. 3c and 4). This matches with previous observations of differential expression of the oxidative phosphorylation pathway in response to mineral weathering^32^. P-type ATPases in fungi are associated with increased transport of K and Na into fungal hyphae^33,34^. Therefore, we attribute the unique expression of the P-type ATPase in +M treatment group to increased transport of K and Na into the fungal hyphae, as observed in our imaging data (Fig. 2a).

**Fig 3.**
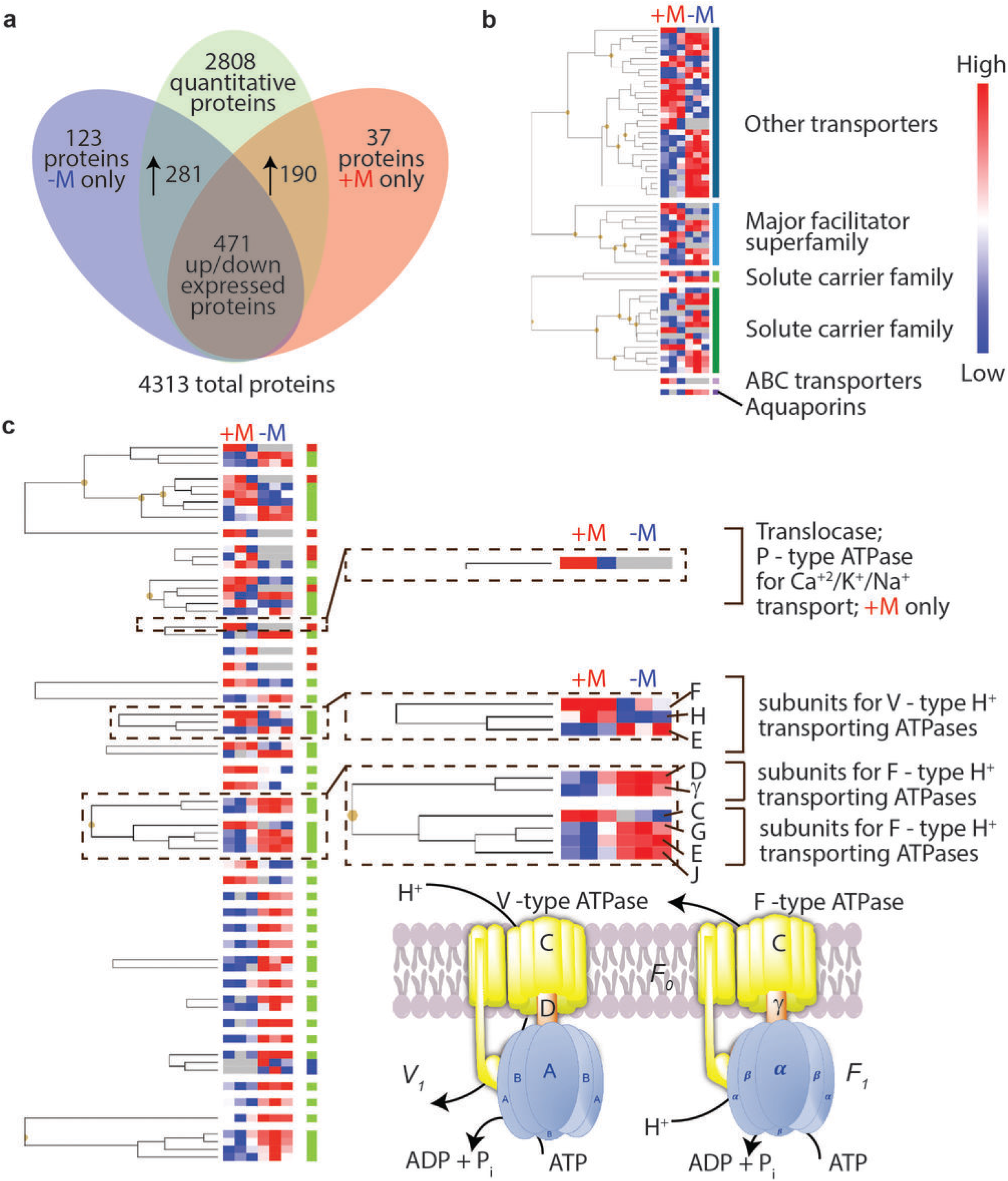
Differentially expressed fungal proteins regulate mineral substrate transport processes. **a**) Venn diagram showing the number of proteins detected and enriched in fungi grown in the presence of minerals (+M) and absence of minerals (-M). **b**) Hierarchical clustering of different transporter superfamilies observed in presence (+M) and absence (-M) of minerals. **c**) Detailed Hierarchical clustering of different transporter types observe in presence (+M) and absence (-M) of minerals. KEGG pathway enrichment analysis demonstrates upregulation of different subunits of membrane ATPases in both treatment groups (+M and -M) related to oxidative phosphorylation pathway (Fig. S12). The P-type ATPase for Ca^+^/K^+^/Na^+^ transport was observed in the +M treatment group alone.

In addition to the unique expression of P-type ATPase for Na and K transport described above, we observed nine unique transmembrane transporter proteins, such as a drug transporter (K03446), electrochemical potential driven transporters (K16261, K07300), a sugar transporter (K08145), and a xenobiotic transporter (K05658) only in the presence of minerals (+M). Moreover, several fungal proteins associated with organic acid transport (K23630, K03448) were enriched only in the +M treatment (Fig. 4).

**Fig. 4.**
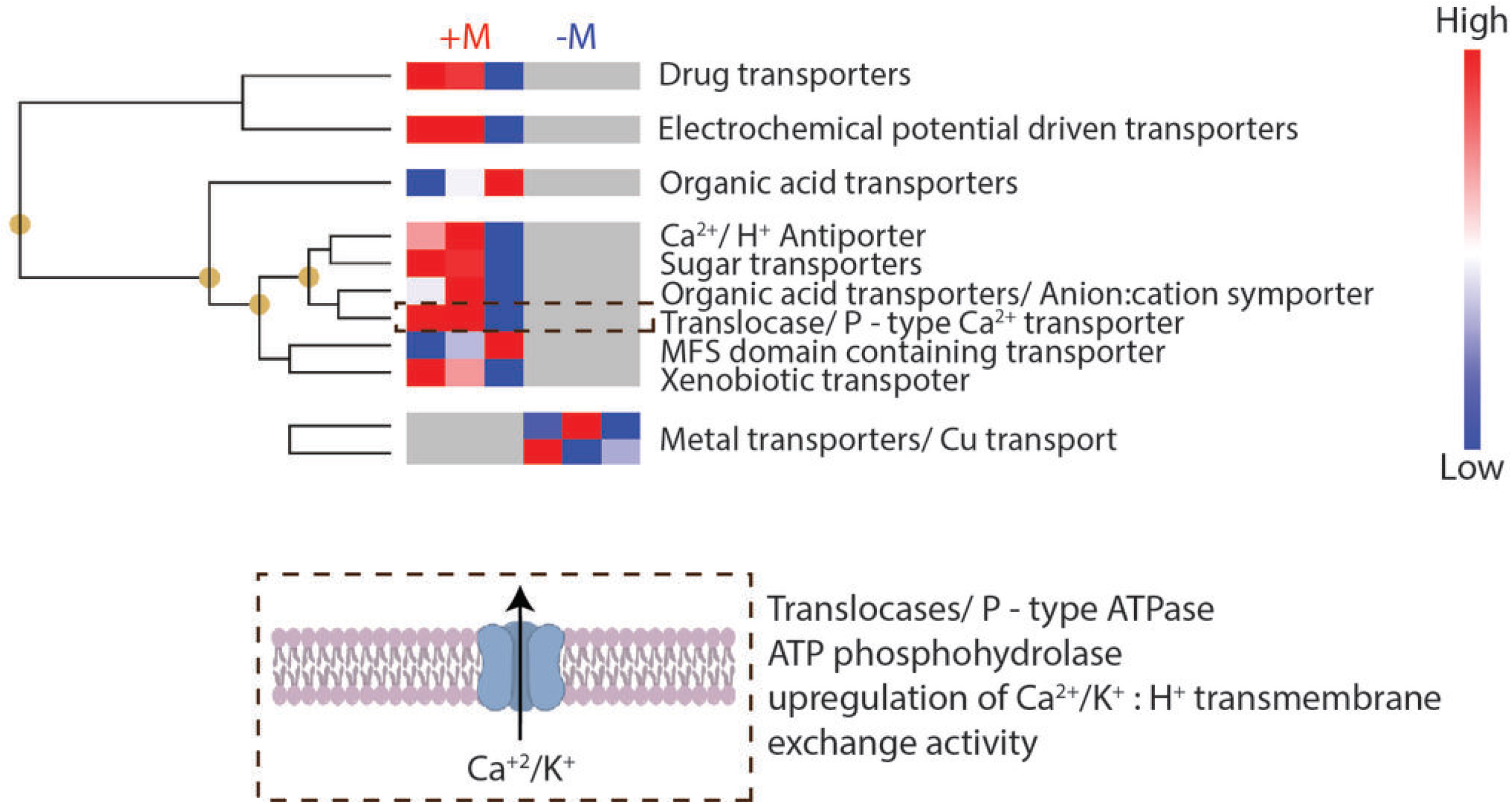
Unique transporter protein expression drives mineral transport in fungi. Hierarchical clustering of different transporter proteins expressed in the presence of minerals (+M) treatment group alone (top). These transporter proteins are not enriched in -M treatment group. The P-type ATPase, part of the oxidative phosphorylation pathway demonstrated in detail in Fig. 3c, was expressed in +M treatment alone. Several organic acid transporters commonly associated with indirect mineral degradation process were also observed only in the +M treatment group. Metal transporters were enriched in -M treatment group, as shown by our analysis. Expression of organic acid transporters in +M treatment group alone suggest *Fusarium sp. DS 682* could use organic acids for extracting nutrients in presence of minerals.

## Discussion

Previous studies on microbial induced mineral weathering have focused on investigating microbial metabolites and secondary minerals that are products of mineral weathering in the bulk soil. However, only a few studies have attempted to decipher weathering mechanisms by exploring the fungi-mineral interface^35^. This is in part due to the spatial heterogeneity of soil and the complexity of mineral weathering processes, which present challenges and necessitate innovative sampling and experimental platforms to study biotic driven mineral weathering. Here, we developed a novel mineral doped soil micromodel platform that emulates the chemical and physical complexity of soil, by embedding solid-phase minerals directly into the polymer matrix to simulate the spatially distinct hotspots of the natural soil environment that drive microbial processes underground. This platform enabled us to visualize *Fusarium sp. DS 682* thigmotropic growth response and mineral dependent increase in hyphal density. We also observed increased K and Na transport and distribution within fungal hyphal networks in presence of minerals (Fig. 2a). K is an essential macronutrient required for fungal growth, while Na uptake by fungi can occur simultaneously or as a substitute for K^36^. Nevertheless, fungal uptake of both Na and K greatly enhances hyphal growth, where hyphal uptake of both nutrients occurs through P-type ATPases^33,34^, as was observed in our proteomics analysis. Here, we observed fungal hyphal bridging between two nutrient plugs (C1 and C2, Fig. 1a and d) only in presence of minerals. Therefore, we propose that the increase in K^+^ and Na^+^ transport promotes hyphal growth to facilitate nutrient foraging in the water and nutrient limitation conditions of the soil micromodel environment.

We also observed changes in the K chemistry as K was transported from minerals into the fungal hyphae, where two distinct types of K chemistries, *inorganic K* and *organic K*, were observed. XANES spectra of the natural kaolinite contained pre- and post-edge features, which are low intensity peaks before and after the highest intensity peaks, which are spectroscopic features consistently seen in other K minerals^37^ (Fig. S7; ‘Powdered mineral’ and ‘Etched mineral’). We observed that the *organic K* spectra in fungal hyphae do not contain pre- and/or post-edge features (Fig. S7; ‘Inoculation point’ and ‘organic complexed’). The absence of these features could be attributed to greater distance or distortion to the nearest-neighbor ions, however more detailed spectroscopy is required to determine the bonding environment of the K in fungal biomass. This result that differentiates *organic* and *inorganic K* spectra is consistent with spectroscopic studies of other mineral systems, where increased crystallinity caused more photoelectron scattering events that generate post-edge features^38,39^. This evidence of changes to the chemistry surrounding K, demonstrates that inorganic K was assimilated by fungi from minerals and suggests potential mechanisms for fungal K mineralization. Our data also present the first description of K XANES of minerals and organic compounds in the context of fungal biology. Further efforts will concentrate on characterizing the transformation and chemistry of mineral K into *organic K* through fungal degradation of minerals.

Collectively, the multimodal chemical imaging revealed that the fungal growth in mineral doped micromodels during nutrient and water limitation occurred through fungal weathering and uptake of mineral derived nutrients. Moreover, we observed increased expression of unique transporter proteins and the presence of organic acid transporters in only the +M condition through fungal proteomics analysis. This evidence supports our hypothesis that fungi are degrading and assimilating mineral derived nutrients to support enhanced growth in mineral doped soil-like environments compared to the mineral free control. Furthermore, the enriched expression of organic acid transporters in the +M condition, together with the absence of micro-tunnels or pores on mineral grains (Fig. S9), suggests that the *Fusarium sp. DS 682* degrades minerals through indirect weathering.

The K transport into fungal hyphae, K speciation in the micromodel minerals, and the presence of organic acid transporter proteins in fungi grown with mineral together suggest that *Fusarium sp. DS 682* has a strong weathering effect on minerals through indirect mechanisms. Previous work on biotic mineral weathering suggests that fungi can sense different mineral types and direct resources toward mineral weathering under specific conditions, such as low moisture and nutrient environments^40,41^. However, the underlying molecular mechanisms by which soil fungi extract and transport mineral-derived nutrients are largely unknown, and direct transport of nutrients to fungi from the mineral surface have not been characterized. Here, by using mineral doped soil micromodels and advanced imaging technologies, we demonstrated mineral weathering and transport of mineral derived nutrients by a saprotrophic soil fungus at the microscale-level of resolution. This platform can enable future studies of different soil fungi processes related to thigmotropic growth response, mineral dependence, nutrient translocation, and beyond. Finally, these results demonstrate the significance of investigating mineral degradation and nutrient transport processes in different classes of soil fungal communities to generate a comprehensive understanding of biotic weathering mechanisms of soil minerals.

## Methods

### X-ray diffraction

X-ray diffraction was performed using a Rigaku Rapid II microbeam diffractometer (Rigaku Corporation, Tokyo, Japan) equipped with a rotating Cr anode (λ = 2.2910 Å) and a large 2D image plate detector. The patterns were collected in transmission mode from a ∼0.5 mm slice of the PDMS/kaolinite composite, and a small aggregate of kaolinite mounted in a glass fiber. A 300 μm diameter incident beam was used, and the image plate intensities converted to powder traces using Rigaku 2DP software. Quantitation of the mineral composition was carried out by whole-pattern (Rietveld) fitting with TOPAS v6 (Bruker AXS) and crystal structures published in the Inorganic Crystal Structure Database (Fachinformationszentrum Karlsruhe, Germany). This method provides the relative fractions of crystalline minerals observed. The natural kaolinite contains relative fractions of kaolinite (85%), K-feldspar (7%), quartz (4%), and mica (4%). The analysis of these fractions is scaled to 100%, ignoring any amorphous materials.

### SEM and EDX analysis

Sections were cut out from PDMS micromodels for SEM analysis to fit into the SEM sample holder. For analysis of PDMS micromodel surface after fungal growth (Fig. S9), fungal biomass was removed by exposing the PDMS surface with fungi to 2% (w/v) N-Dodecyl β-D-maltoside in water. After which, sections were cut from the PDMS micromodel to fit the SEM sample holder. All micromodel sections were mounted into aluminum stubs. To reduce sample charging, the sections were coated with ∼15 nm carbon using a thermal evaporation method (108C Auto Carbon Coater, Ted Pella, Inc.) and were secured with Cu sticky tape on the SEM stage. SEM analyses were conducted with a FEI Helios NanoLab 600i field emission electron microscope. Images were collected with Everhart-Thornley secondary electron detector (ETD) in a field free mode and at acceleration voltage of 3 to 5 kV, current of 0.086 to 0.17 nA and ∼ 4 mm working distance. Backscattering images were also collected, applying low (3 kV) and high (10 kV) settings to identify PMDS locations (hyphal tracks, Fig. S9) occupied by fungal hyphae and correlate the hyphal tracks with positions of mineral grains that were incorporated within PDMS.

The chemical composition of grains in the PDMS and elemental distribution maps, which illustrate hyphae paths and distribution of mineral grains, were collected with an energy dispersive X-ray detector (EDX). The SEM was equipped with an EDX X-Max 80mm^2^ Silicon Drift Detector (Oxford Instruments, Abington, UK) capable of detecting lighter elements. EDX analyses were performed at 3 kV to visualize C-elemental distribution and at 10 kV to collect chemical data on K, Ca, Mg, Si, and Al. Oxford INCA software was used to collect compositional maps (acquisition time > 300 s) and point spectrum analyses (acquisition time ∼80 s).

### Fungal strain extraction and isolation

The soil fungus *Fusarium sp. DS 682* was used for the experiments in this study. This strain was isolated from Kansas Prairie Biological Station (KNZ), as previously described^26^. Fungal spores were obtained from the fungus to use as inoculum for the experiments. To extract spores, 10 ml of 10% sterile glycerol in DI water was added to a hyphal mat of *F. DS 682* grown on potato dextrose agar (PDA) plates for 6 days. The plate was gently tipped in all directions to ensure even distribution of glycerol. The liquid from the plate was removed using a sterile pipette and transferred through a 40 µm filter cap into a 50 ml Falcon tube. Then 1 mL aliquots of the spore suspension were transferred into 1.5 ml safe-lock Eppendorf tubes and stored at -80° C until use.

### Fungal strain culture conditions

100 µl of *Fusarium sp. DS 682* spore suspension were added to M9 PDA plates and incubated at 28° C for 2 weeks. M9 PDA medium was prepared by adding micronutrients (see recipe below), 24 g/L potato dextrose (BD Biosciences, San Jose, CA) and 2% granulated agar (BD Biosciences) to M9 salts (MP Biomedicals, Irvine, CA) in Millipore water. The micronutrients consisted of 0.145 mM ammonium molybdate (Thermo Fisher Scientific), 2 mM boric acid (Sigma Aldrich), 0.151 mM cobalt chloride (Sigma Aldrich), 0.048 mM cupric sulfate (Sigma Aldrich), 0.404 mM manganese chloride (Sigma Aldrich) and 0.048 mM zinc sulfate (Thermo Fisher Scientific). These micronutrients were added after autoclaving the M9 PDA solution, when the solution was ∼ 55° C. After pouring the M9 PDA into petri dishes, the solidified agar plates were wrapped in parafilm and stored at 4° C for future use.

### Micromodel Si master fabrication

The soil micromodels were generated using the SU8 photolithography and PDMS soft lithography processes^42^. Briefly, a chrome mask with the micromodel pattern (Fig. 1) was generated in-house using a mask printer (Intelligent Micro Patterning, LLC, St. Petersburg, FL). The silicon masters were prepared by spin coating SU8 25 (Kayaku Advanced Materials Inc. Westborough, MA) at subsequent spin speeds of 600 and 2000 rpm for 10 and 30 s, respectively, to create a thin SU8 photoresist film using a WS-400B-6NPP/LITE spin coater (Laurell Technologies Corp., North Wales, PA). The silicon wafer with the SU8 layer was heated at 65° C and 95° C for 3 and 7 min for prebaking, respectively, before exposing to 350 nm wavelength of light at 1 W/cm^2^ for 40 s using a mask aligner (NXQ-4006 Contact Photomask Aligner, Neutronix Quintel, Morgan Hill, CA) to crosslink the photoresist. After UV exposure, the Si wafer was baked at 65° C and 95° C for 1 and 4 min, respectively, for postexposure bake and developed using SU8 developer (Kayaku Advanced Materials Inc. Westborough, MA). Finally, the Si wafer with the micromodel image was hard baked at 180° C for 20 min. The micromodel contained pore throats of 12 µm with pillars, representing soil aggregates 300 µm placed 150 µm apart.

### Micromodel fabrication

Polydimethylsiloxane (PDMS; Dow Corning, Midland, MI) soil micromodels were created by mixing the PDMS base and curing polymers at a 10:1 ratio and poured over the Si micromodel channel master. The PDMS was then cured at 70° C for 2 h. To create PDMS micromodels that contained minerals, the same base and curing polymer mixing ratio was used and natural kaolinite powder (Sigma Aldrich, St. Louis, MO) was added at 10% (w/v) to the liquid PDMS and mixed well with a spatula. The PDMS mineral mixture was poured over the SU8 negative channels, and the mineral grains were allowed to sediment overnight (∼ 15 h) and cured at 70° C for 2 h. For reversible binding of the PDMS soil micromodels to glass coverslips, 24 mm × 60 mm glass coverslips (Chemglass Lifesciences, Vineland, NJ) were coated with a layer of uncured PDMS (base to curing polymers ratio of 20:1) by spin coating at subsequent spin steps of 600 and 3000 rpm for 10 and 30 s, respectively. They were then cured by placing the coated coverslip on a hot plate for 10 min at 110° C, creating a sticky PDMS substrate. Creating inoculation port covers for the devices followed the same 20:1 ratio PDMS recipe, and 20 mL of this mixture were poured into a 150 mm diameter petri dish (VWR, Radnor, PA) and cured at 70° C for 30 min. The cured polymer was cut into 6 mm × 6 mm squares and plasma cleaned (PX250, Nordson March, Concord, CA) before using to seal inoculation ports of the micromodels.

### Micromodel etching

The micromodels surface was etched to expose minerals embedded in the PDMS matrix (Fig. S2) using deep reactive ion etching (Plasmalab System 100, Oxford Instruments America Inc., Concord, MA) using a combination of sulfur hexafluoride (45 standard cm^2^/min; SCCM) and O_2_ (5 SCCM) gases for 1 min. The etching process had no measurable effect on the mineral chemistry (Fig. S3). After etching, holes were punched at the two ends of the micromodel channel using a 5 mm hole punch, for placing agar plugs (nutrient) and fungal biomass (starting inoculum) (C1 and C2 in Fig. 1a, respectively). The PDMS micromodels and glass coverslips coated with PDMS were cleaned by rinsing with ethanol, sterile DI water, and then dried using compressed nitrogen, before exposing to oxygen plasma (PX250, Nordson March, Concord, CA) for 30 s. The micromodel and PDMS coated glass coverslips were then left undisturbed in parafilm sealed petri dishes on the counter for 24 h to allow diffusion of methyl groups to the surface of the PDMS that prevented irreversible bonding once the PDMS micromodel and PDMS coated glass coverslips were brought in contact. The channels were tested for leaks, and the sticky PDMS coating prevented any fungal growth outside the channel.

### XPS analysis

XPS data shown in Fig. S3 were acquired using a Physical Electronics Quantera Scanning X-ray Microprobe (Physical Electronics Inc., Chanhassen, MN). This system uses a focused monochromatic Al Kα X-ray (1486.7 eV) source for excitation and a spherical section analyzer. The instrument has a 32-element multichannel detection system. The X-ray beam is incident normal to the sample and the photoelectron detector is at 45° off-normal. High energy resolution spectra were collected using a pass-energy of 69.0 eV with a step size of 0.125 eV. For the Ag 3d_5/2_ line, these conditions produced a FWHM of 0.92 eV ± 0.05 eV. The binding energy (BE) scale is calibrated using ISO 15472 Ed. 2 Surface Chemical Analysis - XPS - Calibration of energy scales. The Cu 2p_3/2_ feature is set at 932.62 ± 0.05 eV and Au 4f_7/2_ line was set at 83.96 ± 0.05 eV. The sample experienced variable degrees of charging. Low energy electrons at ∼1 eV, 20μA and low energy Ar+ ions were used to minimize this charging. Quantification was performed using Ulvac-phi Inc., MultiPak software version 9.1.1.7.

### Fungal inoculation and growth conditions in soil micromodels

A disposable 1.5 mm holepunch (Integra Miltex, York, PA) was used to generate agar pieces to fill into the inoculation port (1 agar plug) and the second nutrient port (4 agar plugs). Using another 1.5 mm holepunch, fungal biomass was collected from a fungal culture agar plate (1 agar plug with fungi) and inoculated in the inoculation port. The agar plugs were generated from PDA without M9 media and micronutrient addition. The ports were then sealed with ∼2 mm thick sticky PDMS inoculation port cover membranes. The micromodels with fungal biomass inoculation were incubated at 28° C for 7 d.

### Characterization of fungal growth and hyphal density analysis

Fungal growth in the micromodels were imaged using a Nikon eclipse TE2000-E epifluorescence microscope (Nikon, Melville, NY). Mosaic brightfield images were obtained using a 4 × air objective and stitched across the length and width of the microchannel to capture fungal growth in micromodels. Prior to processing, each image was cropped to the same square dimensions (6115 pixels x 6115 pixels) to analyze only the area shown in Fig. S5. The structured region was not included in the image analysis, since in micromodels without minerals, fungi did not grow into the pore spaces and the structures created artifacts that convoluted the image analysis results. The cropped images were imported into ImageJ^43^ and each image type was changed to a 8-bit image before further processing. Next, the image threshold was adjusted manually to create the most contrast between the fungal hyphae and the sample background. The image was then inverted, making the fungal biomass appear white against a black background, and the images were made binary, which sets a threshold at where ImageJ determines if a pixel is either black or white (collapsing the 256 intensity channels down to two). The hyphal density was then determined by the ratio of white pixels in an image to the total image. Ten devices per condition were analyzed to obtain the average density and standard deviation per condition.

### Functionalization of PDMS surface

The surface chemistry of the PDMS micromodels without minerals were changed to determine the effect of surface chemistry on fungal growth (Fig. S4b). The surfaces were rendered hydrophilic or hydrophobic by vapor phase functionalization using 3-aminopropyltrimethoxy silane or chlorotrimethylsilane (Sigma Aldrich, St. Louis, MO), respectively. The PDMS micromodels were plasma cleaned for 30 s and placed in a vacuum desiccator (Cole-Parmer, Vernon Hills, IL) along with 100 µl silane solution in an aluminum boat. The micromodels were functionalized after placing in vacuum for 15 h and followed by baking at 70° C for 2 h.

### ToF-SIMS analysis

The micromodels with fungal growth were disassembled by removing the PDMS membrane covers and PDMS coated glass coverslips and were then analyzed using a TOF.SIMS5 instrument (IONTOF GmbH, Münster, Germany). A 25 keV pulsed Bi_3_^+^ beam was focused to ∼5 µm diameter and scanned over a 500 µm × 500 µm area to collect SIMS spectra and images. The current of the pulsed Bi_3_^+^ beam (10 kHz) was 0.56 pA, while the data collection time was about 328 s per spectrum with a mass resolving power between 4000-7000 across the spectra. A low energy (10 eV) electron flood gun was used for charge compensation in all measurements. The ToF-SIMS data was analyzed using SurfaceLab software (Version 6.4), provided by the instrument manufacturer.

### Synchrotron Experiments

The micromodels with fungal growth were disassembled by removing the PDMS membrane covers and PDMS coated glass coverslips. The micromodel surface with fungi was then exposed to 4% paraformaldehyde (PFA) vapors by placing opened micromodels with a PFA-soaked filter in a petri dish for 24 h at room temperature, which arrested fungal growth. Micromodels that contained no fungi, as well as a natural kaolinite powder, were used as controls and these samples were not exposed to the PFA protocol. Micro-X-ray fluorescence (µ-XRF) imaging and X-ray absorption near edge structure (XANES) spectroscopy were conducted using the tender X-ray microprobe beamline 14-3 at the Stanford Synchrotron Radiation Lightsource (SSRL), SLAC National Accelerator Laboratory. Both µ-XRF imaging and XANES spectroscopy were performed around the potassium K-edge (3608 eV) using a water-cooled double Si crystal (111) monochromator to select the incident X-ray energy with a beam size of 5 µm. Natural kaolinite powder (Sigma Aldrich, St. louis, MO) was used to calibrate the monochromator using the position of the white line at 3618.73 eV. The micromodels were mounted onto a 360° rotating sample wheel and placed into the He purged sample chamber (O_2_ in the sample chamber was purged to < 0.02% before analyzing the sample in order to reduce argon interference). Initially, µ-XRF images were collected at a coarse resolution (10 µm step size) above the potassium K-edge (3625 eV) in order to capture all chemical forms of K. Additionally, XANES spectra were obtained on: (i) natural kaolinite powder, (ii) minerals in the micromodel without fungal growth, (iii) minerals in the micromodel with fungi growth, (iv) fungi inoculation point, and (v) a micromodel with fungi growth but without minerals (Fig. S7; defined as end-member spectra) in order to determine the number of different chemical forms of K present in the samples for multi-energy (ME) mapping. ME maps were collected at 3615.65, 3617.25, 3618.4, 3620.15, and 3627 eV based on features identified in the XANES spectroscopy end-members (organic K and kaolinite mineral standards). High spatial resolution ME maps were acquired with a 3 µm step size in order to oversample the regions of interest.

### Synchrotron Data Processing

Data was processed using the MicroAnalysis toolkit^44^, SixPACK, and Athena X-ray absorption spectroscopy software packages (for XANES spectroscopy)^45,46^. During data collection, principal component analysis (PCA) was employed on the ME maps to determine unique components within the mapped region to guide the locations for further investigation with XANES spectroscopy. XANES spectra were processed using a detector deadtime correction, background subtraction of the linearized pre- and post-edge, and edge-step normalization. XANES spectra were fit using a linear combination fitting of the end-member spectra (i.e., the only known K complexes within the sample; Figs. S7 and S8). Subsequently, a cross-plot of the individual components from PCA of the processed XANES spectra from each mapped region was used to identify the spectra that are the most dissimilar to one another (i.e., most different K chemistries). A least-squares fitting of these spectra to the ME maps was generated to a second set of spatial abundance maps of the same region in order to further show the distinct K chemistries^47–49^. Prior to this experiment the differences between XANES spectra of inorganic vs. organic complexed K was unknown. In these experiments, K complexed inorganically (in minerals) and organically (in fungal hyphae) had very similar white line positions, but with different characteristics in the pre- and post-edge. This meant that each ME map appears very similar, and a least-squares XANES fitting was required to tease apart the locations of organic complexed K from mineral-bound K.

### Fungal Proteomics

The fungal samples for proteomics analysis were extracted from fungal biomass grown on PDA agar without M9 and micronutrients. To expose the fungus to minerals, 100 µl of 50% (w/v) natural kaolinite solution in water were autoclaved and spread over the agar surface using a disposable L shaped spreader (Thermo Fisher Scientific, Waltham, MA). The solution was dried for 1 h to create a thin mineral film on the agar surface. Three plates each with (+ mineral) and without mineral (- mineral) condition were inoculated with fungal biomass from 14 d growth on PDA using a 1.5 mm holepunch. The plates were incubated at 28° C for 15 d before extracting hyphae (500 mg) from each plate. The proteins were extracted using the MPLEx protocol as previously described^50^, and further details regarding sample preparation and data analysis is provided below.

### Proteomics sample preparation

The hyphal biomass was extracted in microcentrifuge tubes with beads (Omni International, Kennesaw, GA) and 50 mM NH_4_HCO_3_ (pH 8), and lysed using a bead ruptor elite (Omni International, Kennesaw, GA) tissue lyser by homogenizing twice at 6 rpm for 45 s. The microcentrifuge tube with the cell lysate was placed over a 15 ml falcon tube after a hole was poked in the microcentrifuge tube using a 26-gauge needle, where then the lysate was extracted by centrifuging at 4500 x g for 5 min and the microcentrifuge tube with beads were washed using ammonium bicarbonate solution and centrifuged again. The lysates from the two wash cycles were combined.

A solution of 2:1 CH_3_Cl:CH_3_OH was added at 5 times the volume of the cell lysate and vortexed for 30 s and kept on ice for 5 min and vortexed again for 30 s. The solution was centrifuged at 10000 x g for 10 min at 4° C and the protein interlayer was collected and dissolved in 8 M urea. A BCA protein assay^51^ was performed to determine the amount of protein in each sample before further analysis using the global digestion protocol. The protein interlayer solution in urea was then incubated at 60° C after adding dithiothreitol (Sigma Aldrich, St. Louis, MO) at 5 mM concentration for 30 min. The sample was diluted 10-fold in 50 mM NH_4_HCO_3_ and CaCl_2_ was added to a concentration of 1 mM. Protein digestion was performed by incubating in trypsin (1 µg trypsin/50 µg protein; Thermo Fisher Scientific, Waltham, MA) for 3 h. The samples were desalted using 30 mg/ml C18 polymeric reversed phase column (Strata-X 33u, Phenomenex, Torrance, CA) and analyzed using LCMS.

### Liquid chromatography mass spectrometry of protein digests

A nanoACQUITY ultra performance liquid chromatography (LC) with a 2DLC system (Waters, Milford, MA, USA) was used for separation of protein digests. Buffer A (0.1% formic acid in water) and buffer B (0.1% formic acid in acetonitrile) were used as mobile phases for a gradient separation of 180 min. 5 µl protein digests were automatically loaded onto a C18 reversed phase column prepared in-house (70 cm × 70 µm I.D. with 3 µm Jupiter C18 particle size, room temperature) with 100% buffer A at 5 µl/min. The eluted peptides from the C18 column were analyzed using a Q-Exactive Plus Orbitrap MS (Thermo Scientific, San Jose, CA) for high resolution MS and high-energy collision-induced dissociation tandem MS by electrospray ionization. Samples were analyzed using a 180 min LC-MS/MS method, data acquisition was started 15 min after sample injection. Spectra were collected between 375 to 1,800 m/z at a mass resolution of 180,000 (at m/z 200), following by a maximum ion trap time of 180 ms. Peptides were fragmented using a high-energy collision energy level of 32% and a dynamic exclusion time of 30 s for discriminating against previously analyzed ions.

### Proteogenomic Sequence Database Construction for Proteomics Analysis

*Fusarium sp. DS 682* required additional protein sequence annotation support using constructive proteogenomic database search methods^52–54^. *Fusarium sp. DS 682* was recently sequenced^26,55,56^ and annotated using the recently released genome annotation pipeline Funannotate (v1.7.0; Palmer & Stajich, *GitHub* https://github.com/nextgenusfs/funannotate/tree/v1.7.0), which is specific for fungal and higher order eukaryotic genomic annotation. Functional gene mapping for protein predictions of *Fusarium sp. DS 682* resulted in 19,949 predicted gene product protein sequences, for which 44% of the proteome sequences contained a predicted KEGG ortholog (KO) protein annotation assignment (8,849 total genome KO IDs mapped), leaving 56% of the proteome unannotated. To supplement proteome sequence annotation deficiencies, additional searches were performed using BLASTP to incorporate additional mapped reference protein sequence fragments from sub-selected proteomes of corresponding genomes from the closest model orthologs. The model orthologs used were: *Fusarium graminearum* (<u>GCA_900044135.1</u>; mash^57^ distance=0.21 (k=17, s=100000, as suggested by^58^); 7,548 reference protein annotation transfers from 14,162 proteins), *Fusarium pseudograminearum* (<u>GCA_000303195.2</u>, 36.5 Mb; mash distance=0.21; 7,377 reference protein annotation transfers from 12,448 proteins), and *Fusarium oxysporum f. sp. lycopercisi* (<u>GCA_003315725.1</u>, 36.5 Mb; mash distance=0.14; 11,630 reference protein annotation transfers from 16,646 proteins). These species were selected on the basis they represented a functionally diverse set of phylogenetic clades and mimic a similar geographical host/substrate grassland soil environment^59–61^. Ortholog supplementation permitted annotation of an additional ∼16% of the *Fusarium sp. DS 682* proteome.

### Proteogenomic Data Analysis

Label-free *Fusarium sp*. DS 682 raw sequence data files were processed using MaxQuant^62,63^ (v 1.6.7.0) for feature detection, comparative peptide sequence database searches, and protein quantification for subsequent downstream proteomic analysis for biological interpretation. Proteomic sample collection dataset groups for *Fusarium sp*. DS 682 consisted of two fungal growth treatments, with (+M) and without (-M) minerals, where each treatment group contained three biological replicates. Raw fungal dataset files were grouped by assigned treatment group and analyzed using a standard LC-MS run type. N-terminal protein acetylation and methionine oxidation were selectively applied as a variable modification for group-specific parameters, allowing a maximum of three modification per peptide. Peptides used for parent sequence matches required a minimal of one peptide observation per protein, per organism collection file, and a minimum peptide length of seven amino acids. Sequence digest parameters were restricted to an allowed maximum of two missed tryptic cleavages with an MS/MS tolerance of 20 ppm. For increased peptide/protein identification, match between runs (MBR) employed with a 20 min alignment window (0.7 min time match window) using a peptide/protein (unique + razor matches) match false discovery rate cutoff of ≤ 0.01 (1% FDR).

Peptide fragments were quantitatively searched against a constructed proteogenomic sequence database using a suite of organism protein collection files for *F. graminearum* (GIBZE)—<u>Fusarium_graminearum_PH-1_SPROT_TrEMBL_2019-10-16.fasta</u> (downloaded on 10/16/2019 containing 14,160 protein entries), *F. pseudograminearum* (FUSPC)—<u>Fusarium_pseudograminearum_CS3096_SPROT_TrEMBL_2019-12-13.fasta</u> (downloaded on 10/16/2019 containing 12,448 protein entries), *F. oxysporum* (FUSOX) top strains—<u>F_oxysporum_SPROT_TrEMBL_2019-11-21.fasta</u> (downloaded on 10/16/2019 containing 70,763 protein entries), and resulting predicted protein sequence file for *Fusarium sp. DS 682* (SF-1-001.genemark_KOfam_V2_May2020_new.fasta). All other parameters applied not listed here were ran using MaxQuant software defaults.

### Statistical Preprocessing of proteomics data

Statistical pre-processing was performed on resulting dataset MaxQuant output file (‘proteinGroups.txt’) containing iBAQ absolute protein abundances data for both mineral treatment groups and corresponding biological sample replicates (n=3). Datasets were first filtered for removal of all potential contaminants and reverse hits. The following criteria was used to further filter reference sequence database match quality using proteogenomic search methods prior to statistical analysis. Dataset peptide/protein sequence matches were filtered out from the data for either protein groups where 1) the majority protein identifier was comprised of only ortholog sequence file matches, or 2) the majority protein identifier listed had multiple proteins associated with the same redundant peptide fragment matched within a given reference organism collection file. Finally, for assessing reproducibility, mineral treatment groups with too few replicate observations to conduct a quantitative or qualitative statistical comparison were removed (i.e., at least two observed abundance values per mineral treatment group or at least three observed abundance values in one group). The remaining 2,808 proteins passing match quality filter criteria for subsequent statistical analysis were log2 transformed and all non-observed values were assigned a value of NA. Filtering did not change the abundance profiles distributions.

### Statistical Analysis of Proteomics Data

SPANS software^64^ was utilized for evaluating potential data normalization strategies resulting in data normalized via median centering. Differential analysis of log_2_ transformed relative abundance expression data was applied for assessing quantitative (ANOVA) and qualitative (g-test) statistical significance. A one-way analysis of variance (ANOVA, p-value cutoff of 0.05) test was run for each fungal mineral treatment group for comparing mean abundances (log_2_) across biological replicates. A g-test was run for evaluating qualitative differences in the presence/absence patterns of significantly expressed proteins identified in either treatment group using a null hypothesis that presence/absence patterns are not related to a biological group. Resulting ANOVA test flags (quantitative change) and g-test flags (qualitative change) indicating the direction and range of significant change in protein expression values (0: not significantly expressed, 1: significantly expressed in (+M) treatment group, -1: significantly expressed in (-M) treatment group), were filtered (cutoff of +/- 1, ANOVA/g- test flags; ≤ 0.05, ANOVA p-values) for representing the top-most proteins observed to be significantly changing in expression or uniquely observed by presence/absence in one mineral treatment group over the other. 160 proteins were uniquely observed by g-test (presence/absence) in only one mineral protein group. Unique protein observations are defined as being observed in only one mineral treatment group and has abundance values for 2/3 across proteomic sample replicates within a given treatment group. To explore sequence data match quality statistics supplement to this proteogenomic method analysis, see publication data DOI: <u>10.25584/KSOmicsFspDS682/1766303</u> for download.

### Functional Metabolic Pathway Analysis

*Fusarium sp. DS 682* predicted protein KEGG Ortholog (KO) assignments from resulting statistical analyses for quantitative and qualitative significance, for both mineral groups, were mapped to metabolic pathways using KO reference database resource <u>KEGG Mapper</u>.^65,66^ This permitted linking genomes to pathways by functional orthologs. Statistically significant *Fusarium sp. DS 682* proteins not assigned to a predicted KO ID were omitted from KEGG Mapper entity list. However, if an unassigned predicted *Fusarium sp. DS 682* protein entity was matched to a corresponding cross-reference ortholog KO assignment using the MaxQuant database search methods (passing quality match filter criteria above), then the ortholog KO was adopted by default, or an available KO number was obtained using protein sequence BLASTP search results if applicable. KEGG Mapper metabolic outputs linked proteogenomic sequence predictions to functional ortholog annotations for metabolic pathway maps (ortholog molecular interactions, reactions, and relations), BRITE hierarchies (ortholog functional hierarchies of biological pathway or reaction entities), and KEGG modules (complete ortholog modular units of a conserved function, such as conserved reaction transport complexes). 495 of 631 statistically significant proteins observed, matching to a non-redundant predicted KO assignment (iPath_3_ map <u>Fusarium</u> sp. DS 682 Fungal Mineral KO Metabolic Pathways), were mapped to 300 metabolic pathways, 41 BRITE hierarchies, and 105 KEGG modules. Pathway analysis revealed specific metabolic shifts in fungal Central Energy Metabolism in response to mineral addition (iPath_3_ map <u>Fusarium sp. DS 682 Fungal Mineral KO Metabolic Pathway Modules</u>). For mapping predicted transporter protein activity for both treatments, protein KO entities with an observed increase in quantitative abundance (ANOVA, p-value cutoff of 0.05) or unique presence/absence (by g-test = +1) were further evaluated. *Fusarium sp. DS 682* fungal transporter proteins were mapped to 54 metabolic pathway maps, 15 BRITE hierarchies, and 5 KEGG reaction modules, revealing a significant shift in central metabolic energy transfer storage and translocation mechanisms (iPath_3_ map <u>Fusarium sp. DS 682 Fungal Mineral Metabolic Pathway Module Transport</u>). 60 predicted cross-ortholog protein transporters were identified for Fusarium sp. DS 682 identified belonging to 5 major active membrane trafficking involved transporter families (Fig. 3b and S10).

## Data Availability

All data have been deposited at the PNNL <u>DataHUB</u> repository and are available for download under the project data DOI accession: <u>10.25584/KSOmicsFspDS682/1766303</u>. The contents of the data package reported here are the first version and contain required raw and post-processed data for all data acquisition. A comprehensive collection of metadata standard information, a package content “Read Me” file, and ontology data dictionary have been provided at the data package download in compliance with reported guidelines provided by the journal in addition to community standard initiatives supporting FAIR data principles. For increased data availability, fungal isolate *Fusarium sp. DS 682* raw unprocessed mass spectrometry data have been deposited at the MassIVE database repository under the accession <u>MSV000087221</u> and is accessible at ftp://massive.ucsd.edu/MSV000087221/.

## Acknowledgements

This research was supported by the U.S. Department of Energy (DOE) Office of Biological and Environmental Research (BER) and is a contribution of the Scientific Focus Area “Phenotypic response of the soil microbiome to environmental perturbations.” Pacific Northwest National Laboratory (PNNL) is operated for the DOE by Battelle Memorial Institute under Contract DE-AC05-76RLO1830. A portion of the research was performed using the Environmental Molecular Sciences Laboratory, a DOE Office of Science User Facility sponsored by the BER and located at PNNL. Use of the Stanford Synchrotron Radiation Lightsource (SSRL), SLAC National Accelerator Laboratory, is supported by the DOE, Office of Science, Office of Basic Energy Sciences under Contract No. DE-AC02-76SF00515. The SSRL Structural Molecular Biology Program is supported by the DOE-BER, and by the National Institutes of Health (NIH), National Institute of General Medical Sciences (NIGMS, P30GM133894). The contents of this publication are solely the responsibility of the authors and do not necessarily represent the official views of NIGMS or NIH.

## Contributions

AB and CRA designed the experiments and wrote the paper. AB performed all the experiments. OQ performed SEM analysis. JR performed the XANES and XRF analysis. LNA performed proteomics data analysis, created metadata standard information content, prepared data download content for all data files and metadata, created final DOI package. KS helped in creating the soil micromodels. LMB did the proteomics statistical analysis. GL helped maintain fungal cultures. DO performed mass spectrometry analysis on proteomics samples. ZZ performed ToF-SIMS analysis. ME performed XPS analysis. MB performed XRD analysis. WN created the proteomics FASTA file with KO and GO annotations. AJ provided the fungal strains and advised on maintaining fungal cultures. All authors edited the manuscript, with AB, CRA, JJ, KH provided the main edits.

## Figure Legends

**Fig. S1.**
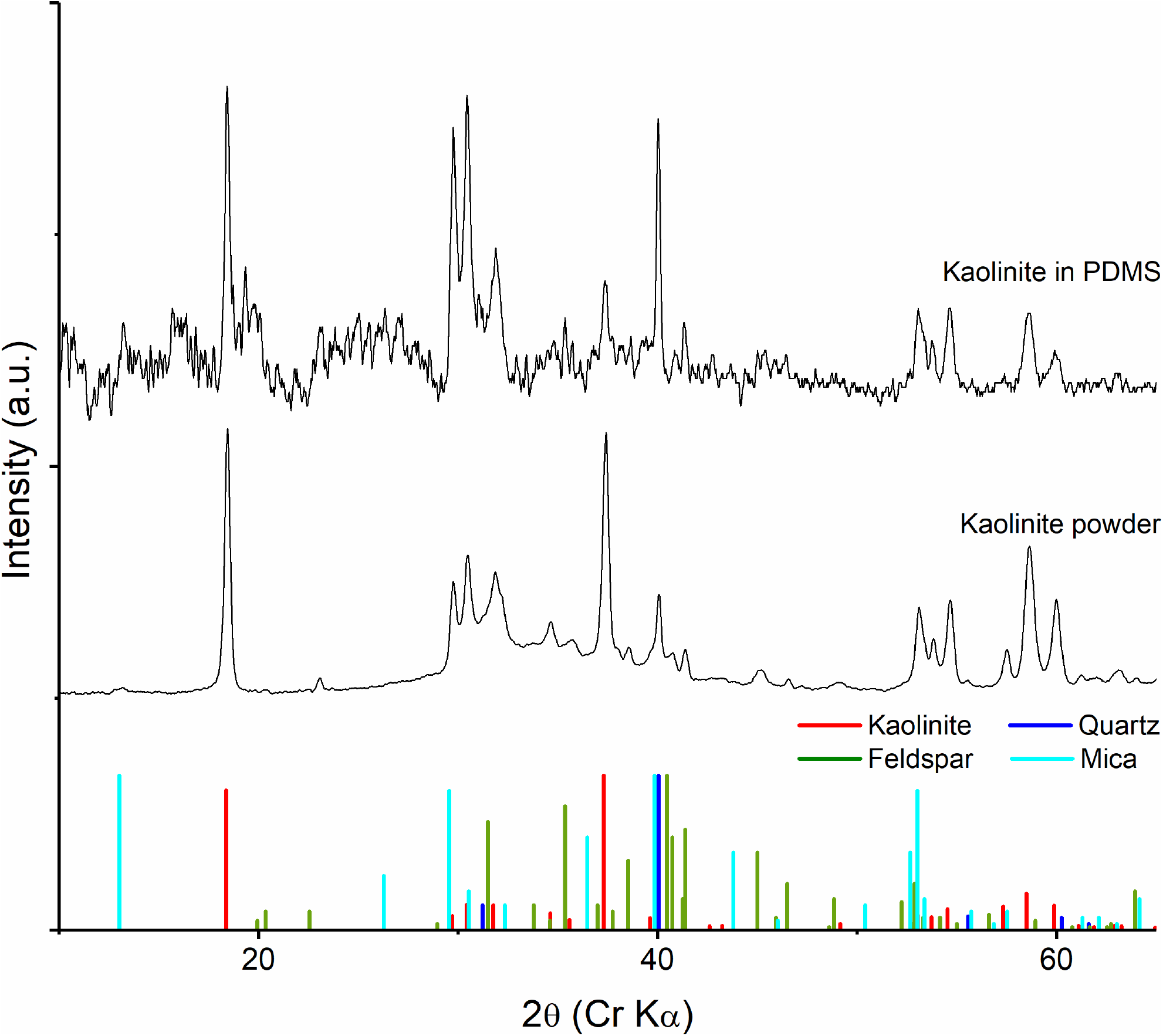
The natural kaolinite powder contained minerals with essential nutrients, which were still present once embedded in the micromodel. X-ray diffraction (XRD) analysis comparison of the natural kaolinite powder and it embedded in the soil micromodels demonstrates the presence of minerals such as feldspar, mica, and quartz, along with kaolinite. The results also show that the mineral doping process within the PDMS does not alter presence of the mineral phases of the natural kaolinite powder.

**Fig. S2.**
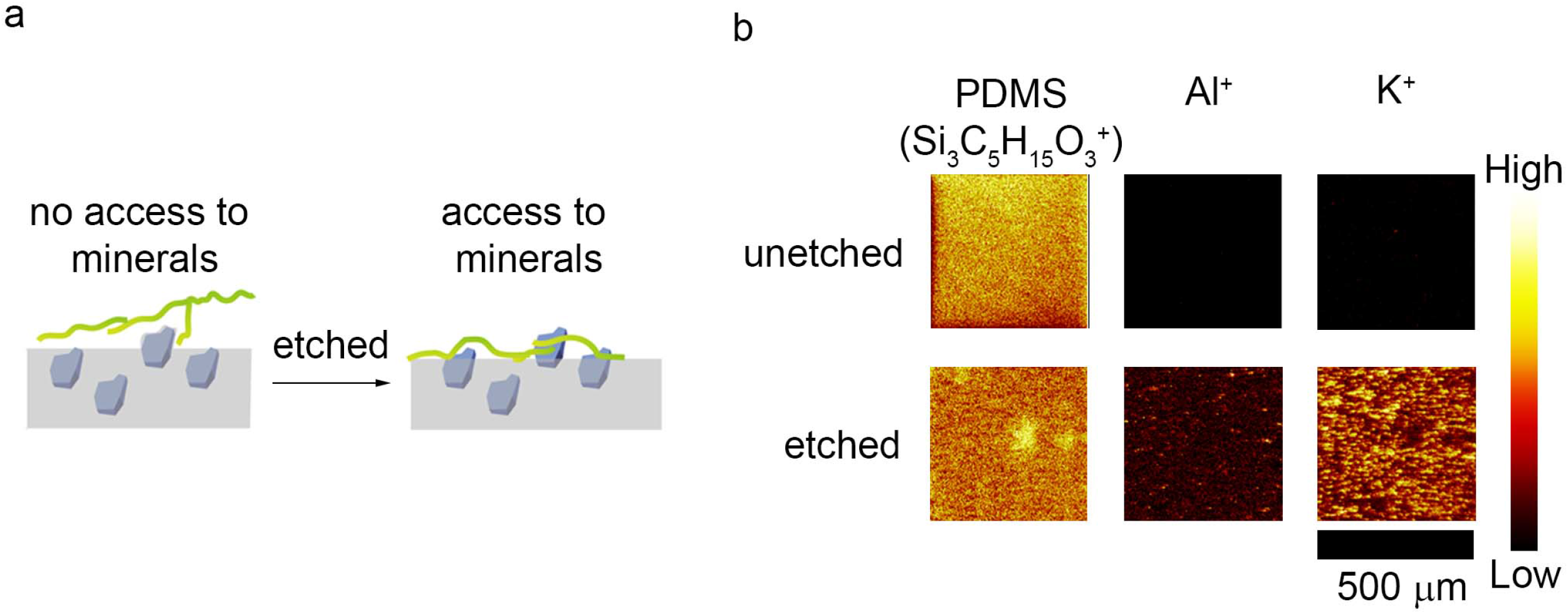
Use of dry etching process helps facilitate access of minerals for fungi on the surface of the micromodel. **a)** A schematic demonstrating that minerals embedded in the PDMS matrix are made available at the surface of the micromodel, for access to fungal mycelia, through deep reactive ion etching (DRIE). We observed fungal growth is limited when minerals are not made available at the surface. **b)** Secondary ion images of the mineral doped micromodel surface before (top row images) and after DRIE (bottom row images). The surface analysis of the micromodels before and after etching demonstrates that K and Al is more available on the surface after etching.

**Fig. S3.**
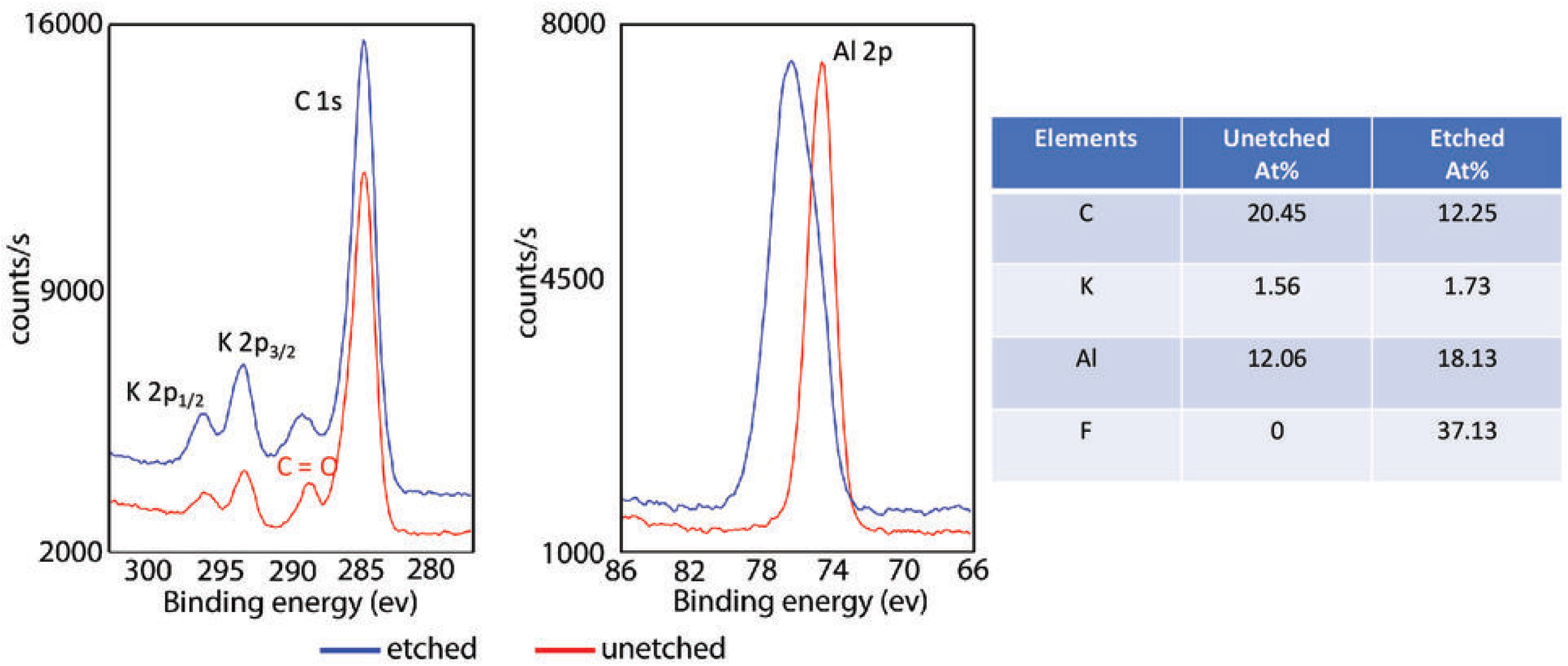
The DRIE etching does not change K coordination chemistry in the minerals. X-ray photoelectron spectroscopy analysis of mineral grains etched by the DRIE process shows binding energy peak shifts of Al 2p corresponding to the formation of AlF_3_. However, peak shifts related to K (i.e., K 2p_1/2_ and K 2p_3/2_) were not observed, indicating the K in the mineral grains remain unchanged after the etching process. These results suggest any observed mineral speciation will be from biotic mineral degradation by the fungi and not from abiotic micromodel fabrication processes. The table on the right shows atomic % of different elements in etched and unetched mineral grains, where fluorine content is increased in etched mineral due to etching by SF_6_ plasma. The K content is comparable in both etched and unetched minerals.

**Fig. S4.**
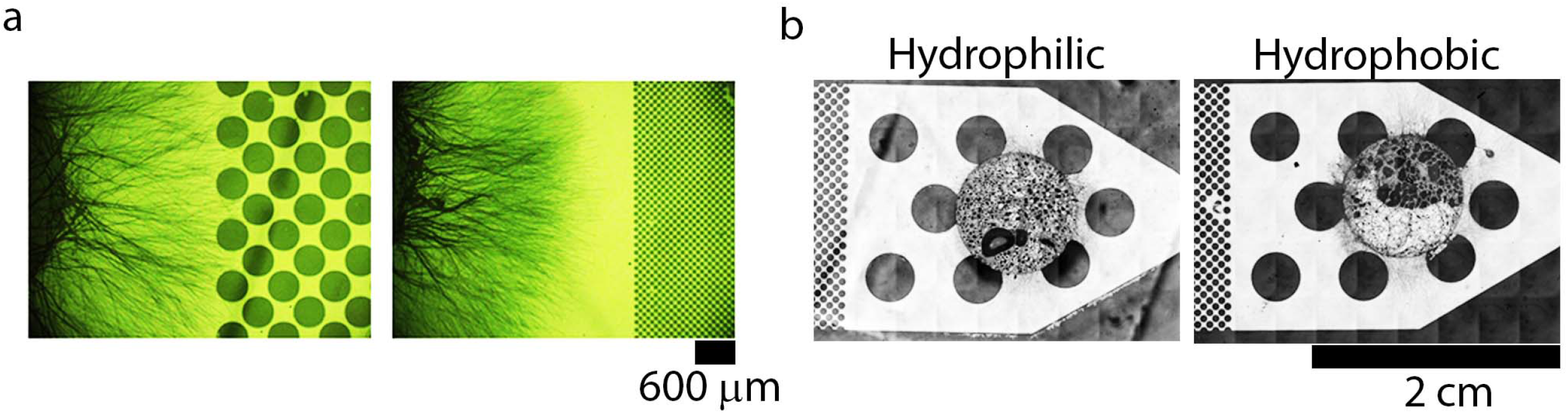
Fungal hyphae growth is dependent on pore size within micromodels, but not the surface chemistry of the PDMS. **a)** Optical images of fungal growth in micromodels with different porosity shows hyphal growth can occur through bigger pore sizes (600 µm), but growth is attenuated in pore spaces closer to those observed in soil (5 µm). **b)** Fungal hyphal growth is not dependent on the surface chemistry of soil micromodels, as shown by the similar growth behavior that was observed in micromodel with PDMS interfaces that were hydrophilic and hydrophobic.

**Fig. S5.**
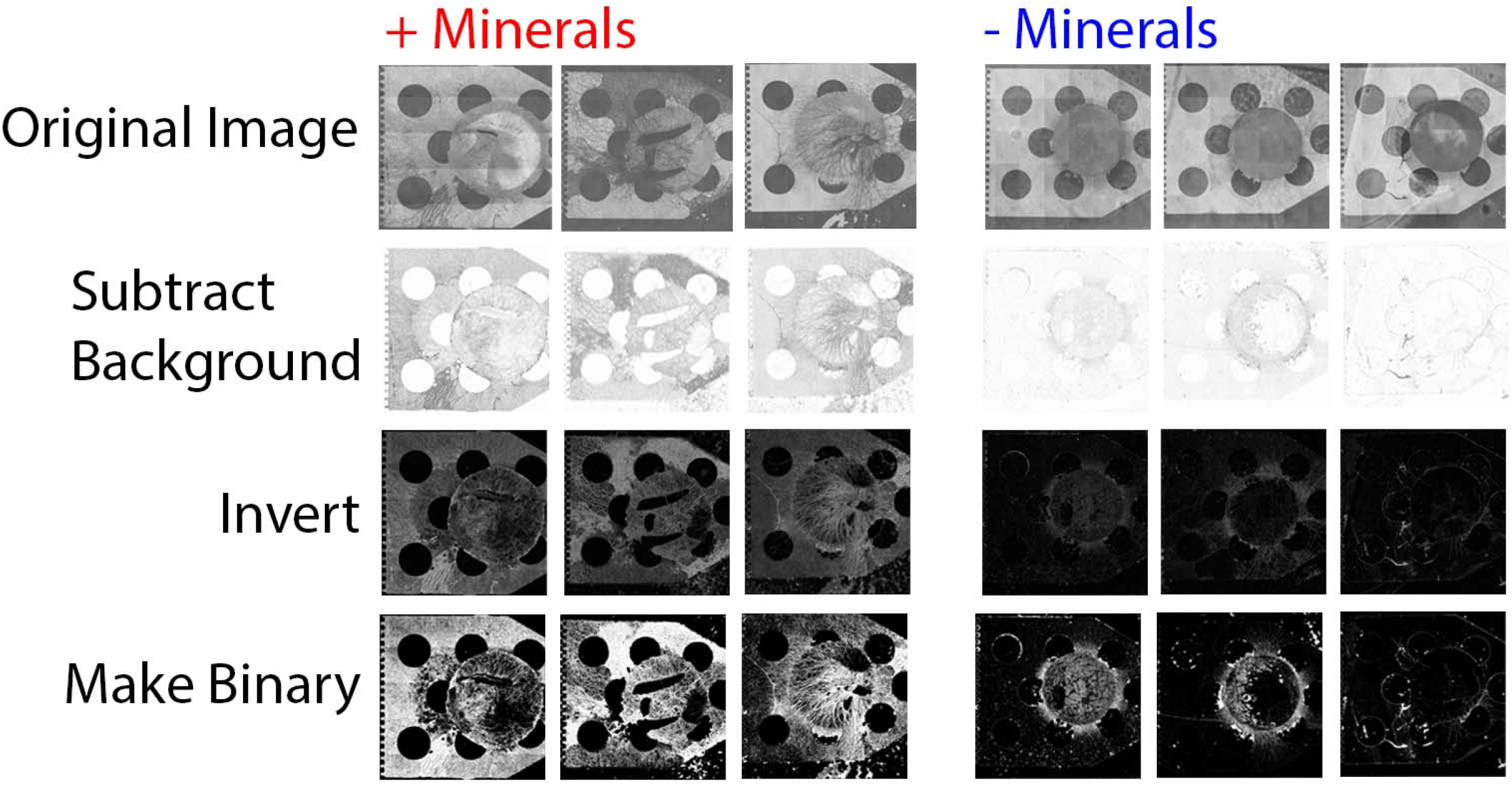
Image analysis workflow demonstrates differences in hyphal density in micromodels with and without minerals. The images demonstrate the workflow using ImageJ software to quantify the hyphal density (Fig. 1d) from optical images. Ten images were used per condition, and we show three different images per condition here to demonstrate the progression of the image analysis to obtain hyphal density of fungal growth with and without minerals.

**Fig. S6.**
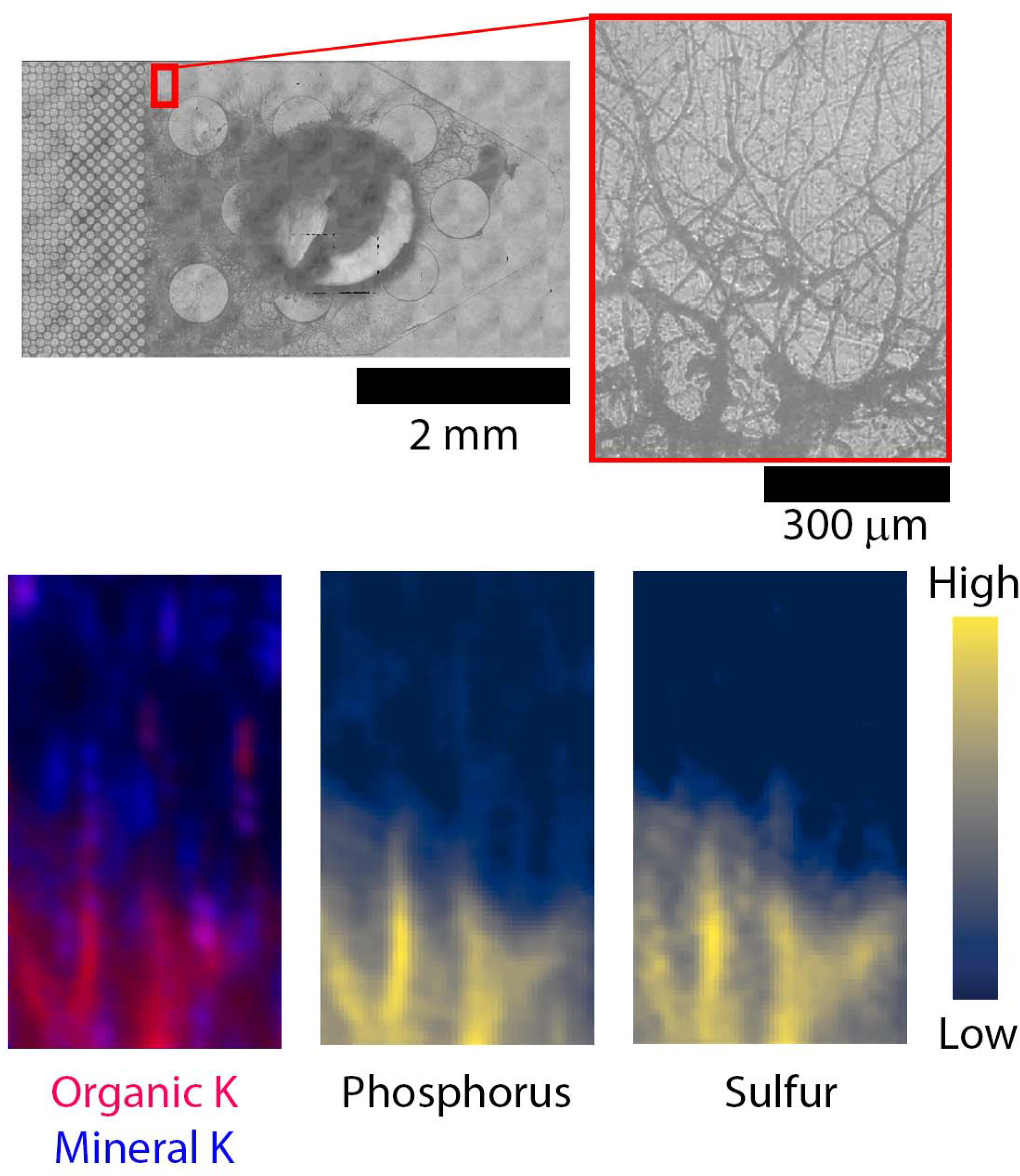
Location of fungal hyphal growth was identified in µ-XRF maps by visualizing the location of P and S. Optical images (top) of µ-XRF mapping region, with specific region notated by the red box and the associated zoomed in image, and µ-XRF maps of organic associated K and mineral associated K (bottom, left), P (bottom, center), and S (bottom, right). Detection of P and S is associated with fungal hyphal biomass in the micromodels, as shown in the optical image of the same region as the XRF maps.

**Fig. S7.**
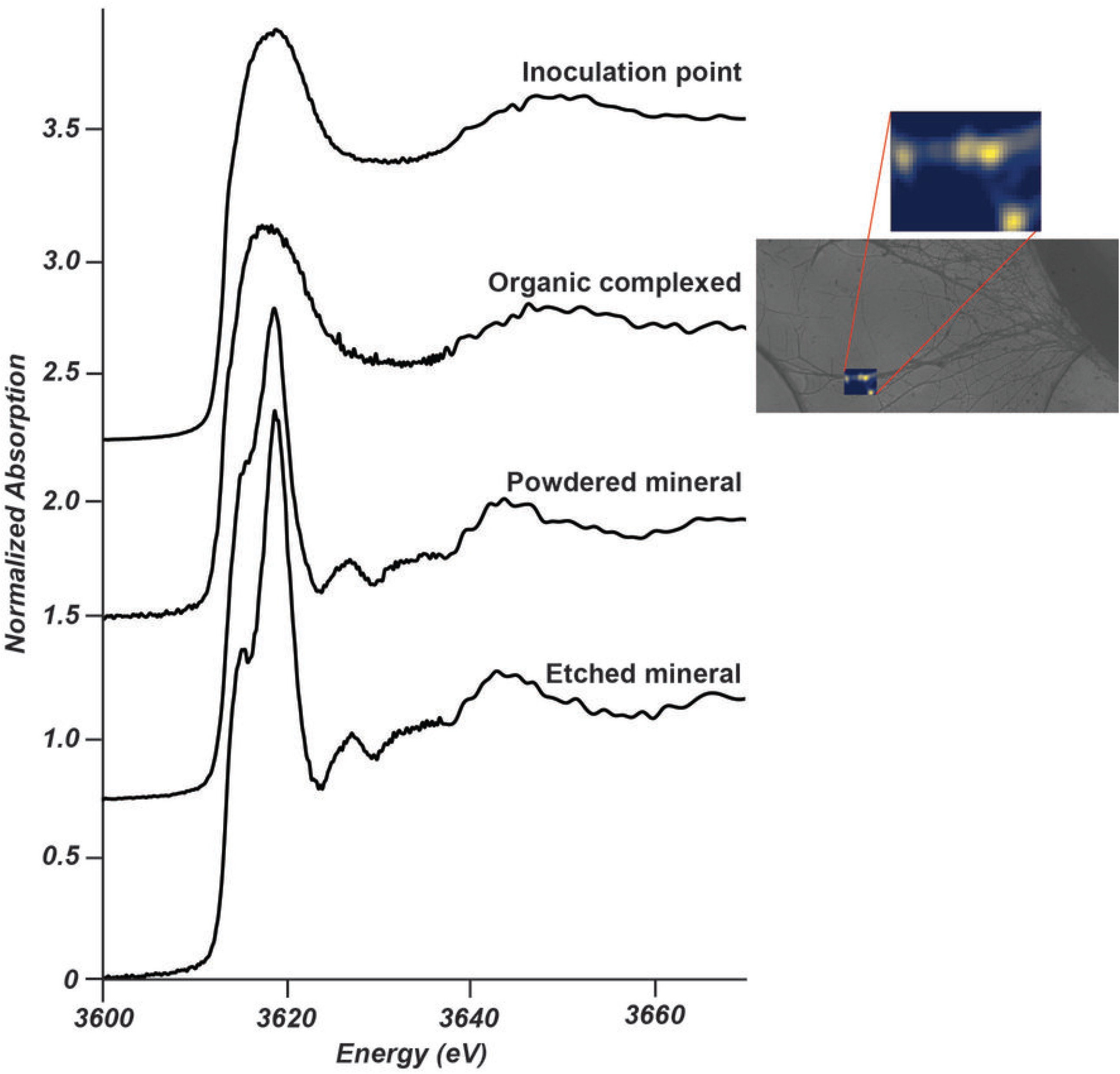
The K bonding environment vary widely between organic- and mineral-derived K. XANES spectra of the different end member controls (solid lines, top to bottom): fungal growth at point of inoculation within mineral doped micromodels, organic complexed K in fungal hyphae in micromodels without minerals, natural kaolinite powder, and in a controlled etched mineral doped micromodel not used for fungal growth. The optical microscopy image next to the organic complexed K spectra shows the fungal hyphae from where XRF maps and XANES spectra of hyphae was generated. Since this represents the first XANES spectral analysis of organic-bound K, we utilized mineral standards (natural kaolinite) and organic K spectra (from fungal inoculation point, C1 in Fig.1a) to model our XANES data using linear combination fitting shown in Fig. S8.

**Fig. S8.**
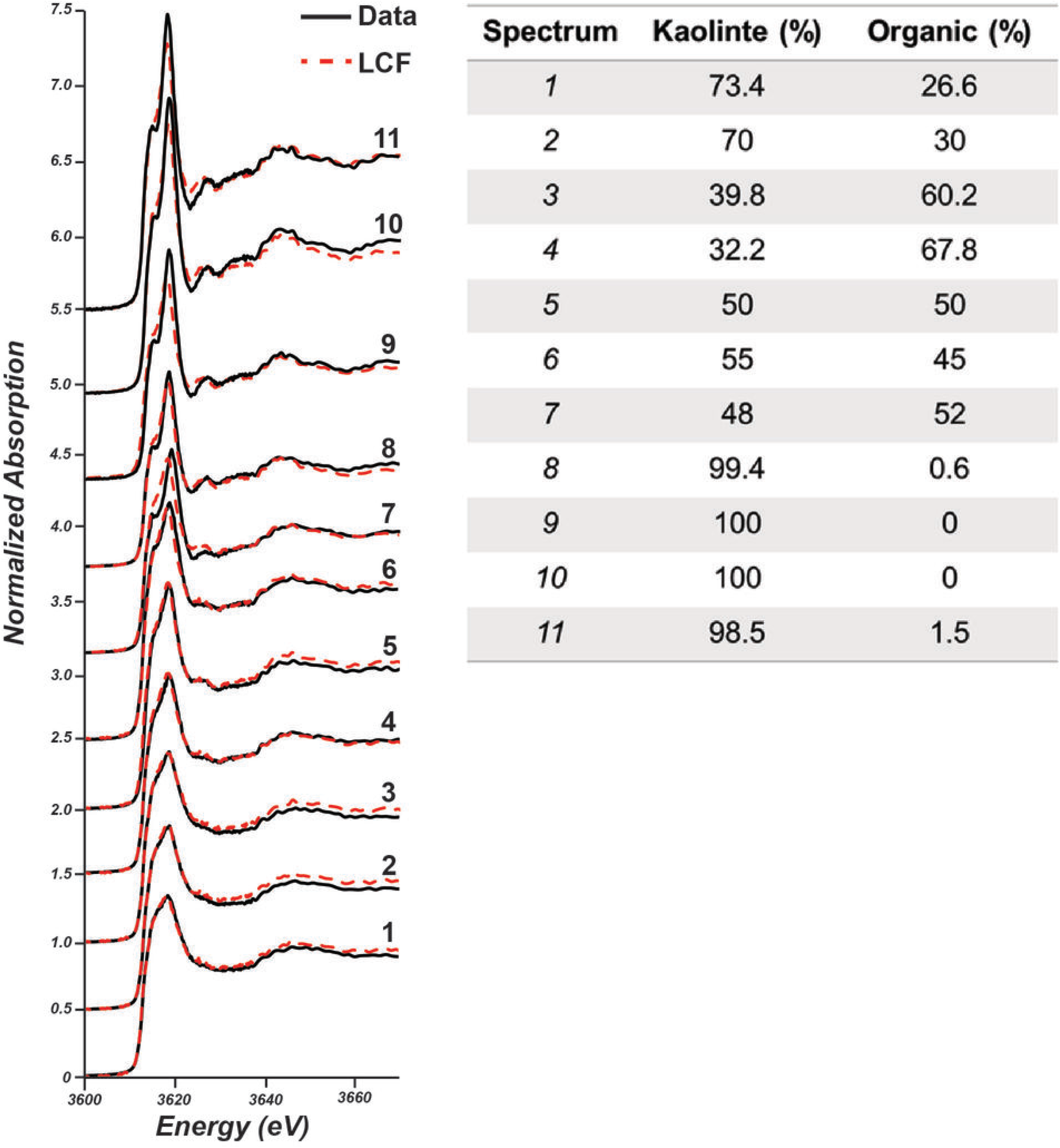
K speciation occurs along with fungal growth in mineral doped micromodel. The linear combination fitting of the XANES spectra from the locations numbered in Fig. 2c. Spectra show a change from organic K (numbers 1 to 6) dominated to mineral bound K (numbers 7 to 11) across the mapped region (left). The linear combination fitting using organic complexed spectra shows different percentages of mineral and organic complexed K on the micromodel corresponding to fungal hyphae and speciated minerals in soil micromodels (right).

**Fig. S9.**
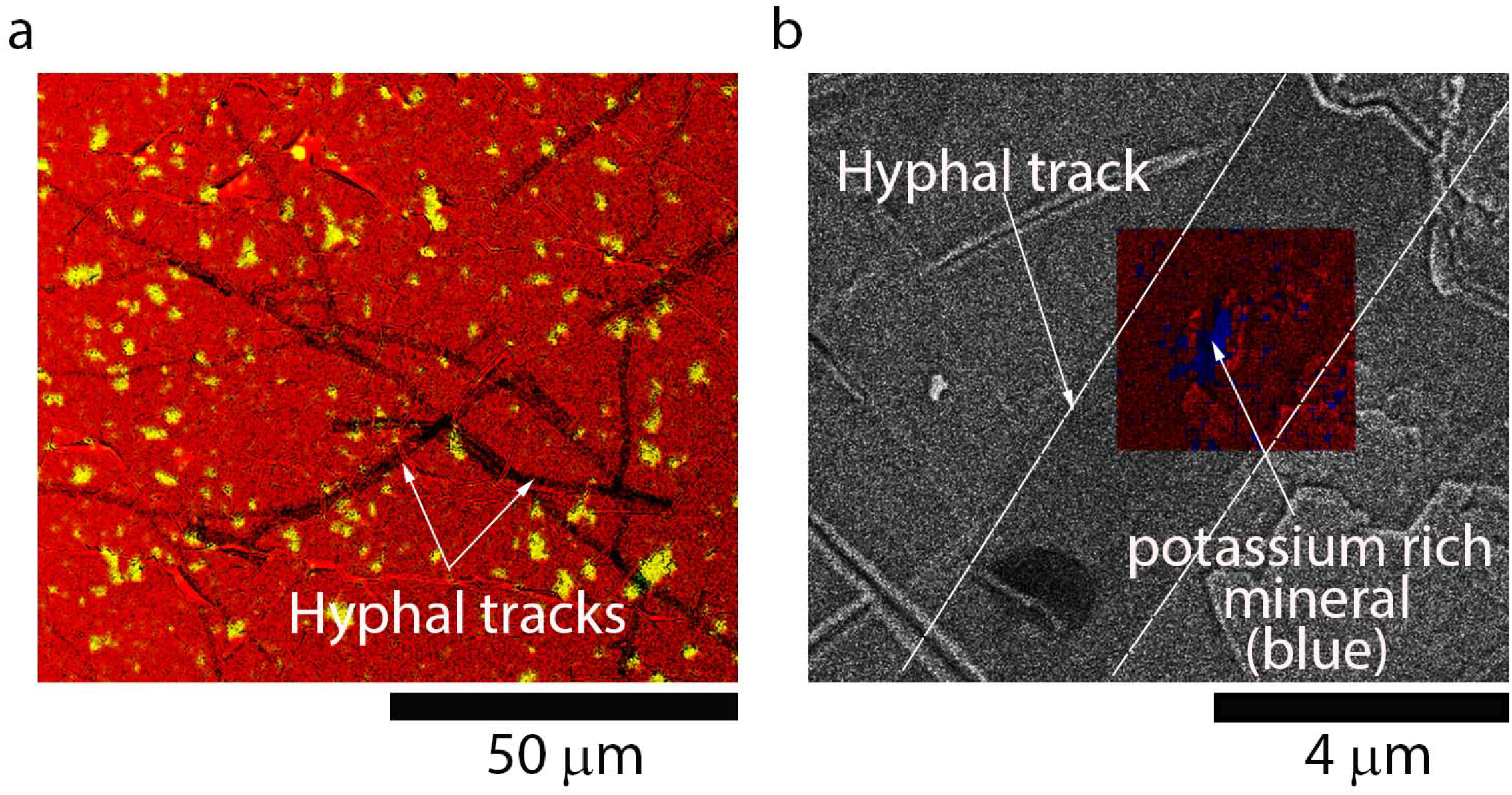
Fungal hyphae track K-containing mineral grains on the micromodel surface. Scanning electron microscopy coupled with energy dispersive X-ray analysis of the micromodel surface after removing fungal biomass shows fungal hyphal tracks on mineral doped micromodel surface. **a)** shows abundance of hyphal tracks, where Al in kaolinite is shown in yellow and **b)** shows a closer look of a hyphal track over a K-rich grain. Hyphal growth was more abundant over K-rich mineral grains as shown in b.

**Fig. S10.**
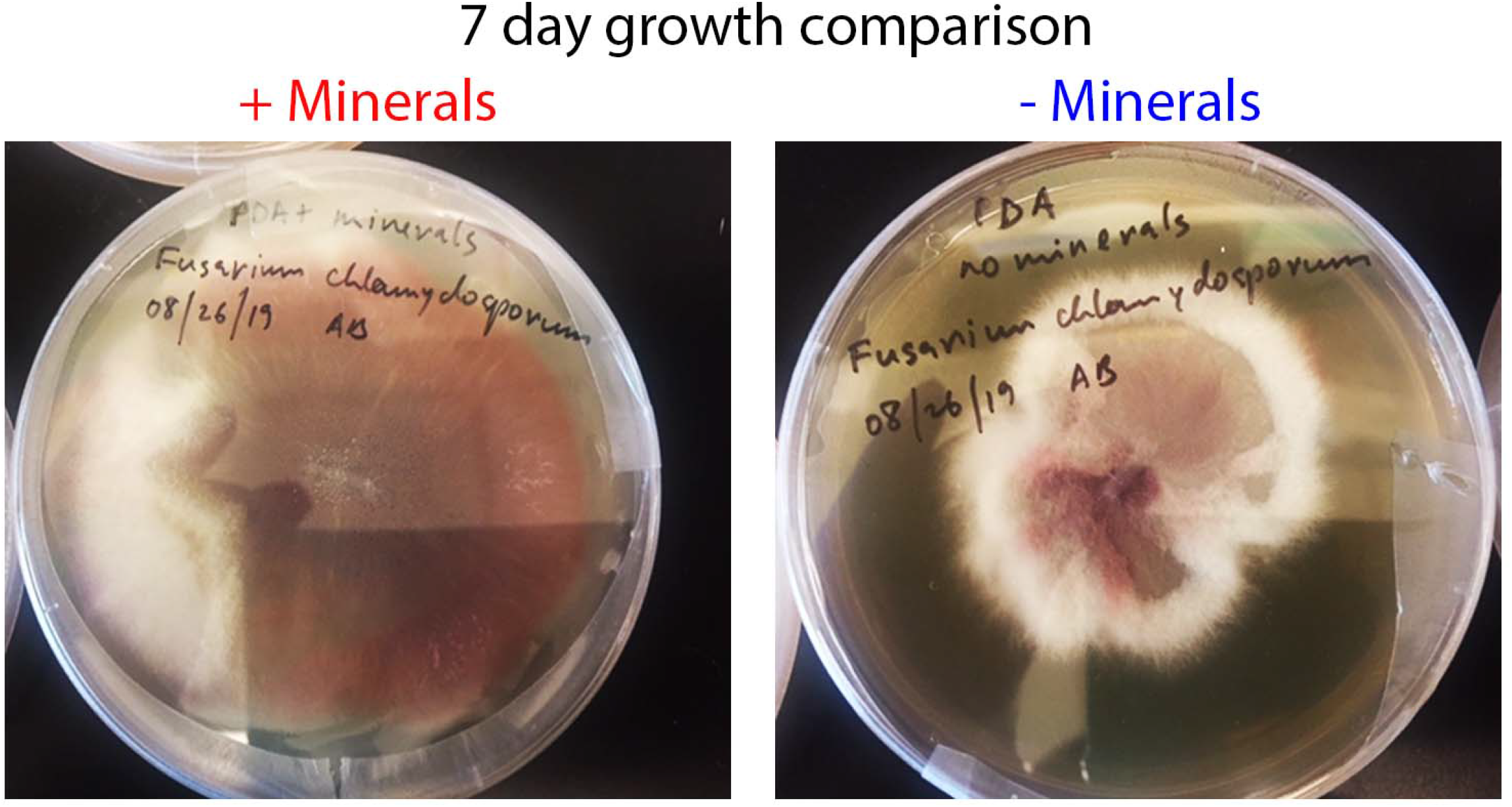
The presence of minerals induces increased fungal growth over time. Fungal biomass grown on potato dextrose agar with (+ Minerals) and without minerals (- Minerals) for 7 days. Faster growth of fungi was observed in the presence of minerals. Since proteomics analysis required a greater amount of fungal biomass than can be harvested from the microfluidic channels of the micromodels, we used the fungal biomass from agar plates amended with and without natural kaolinite. Fungal biomass for proteomics analysis was collected from agar plates as shown here.

**Fig. S11.**
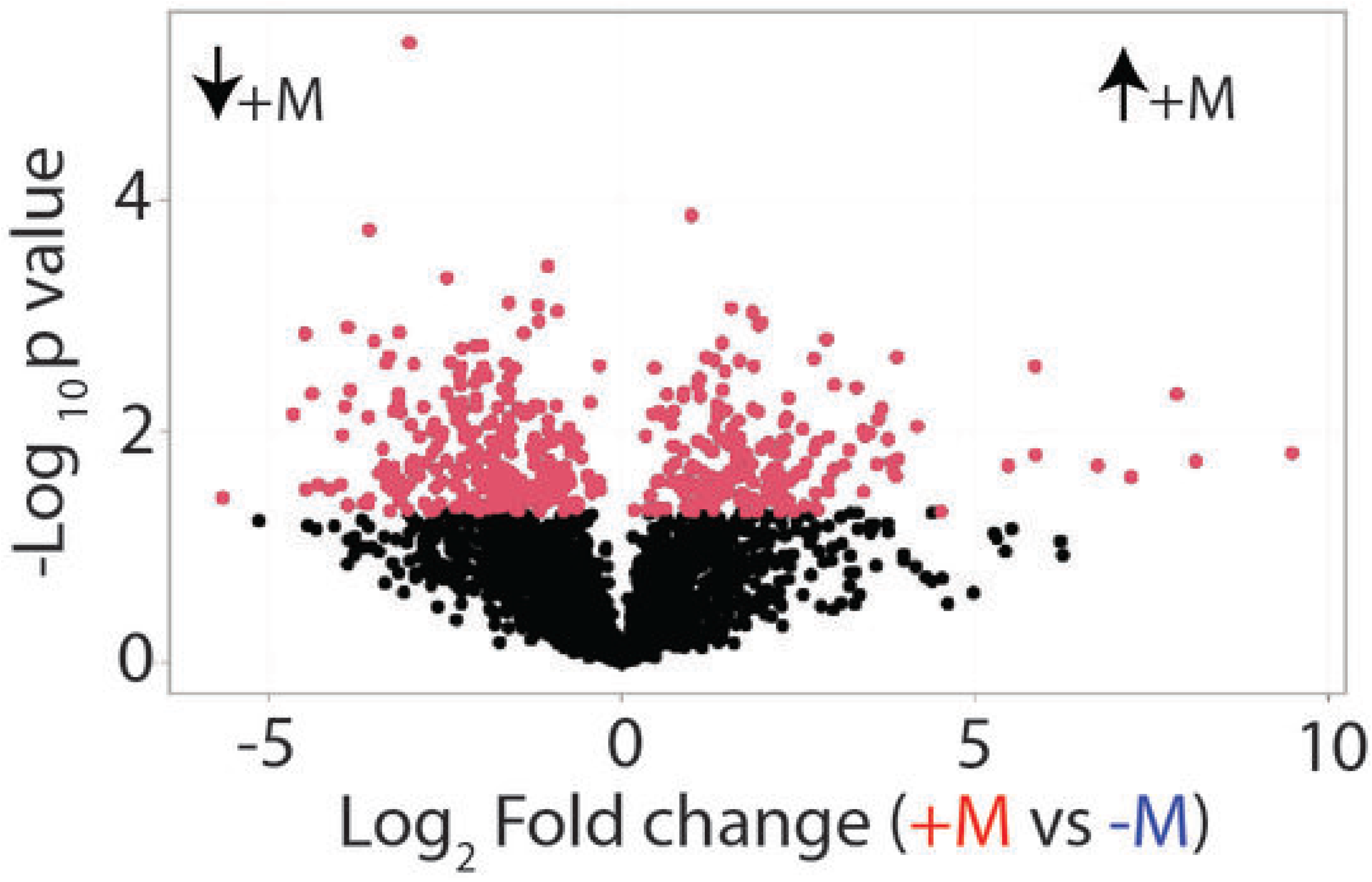
Clear differences are observed in the proteome of *Fusarium DS 682* grown in the presence of minerals compared to growth without minerals supplemented. Volcano plot from ANOVA of proteomics data demonstrate the overall log fold change in protein expression in +M vs -M treatment. Statistically significant protein groups (p-value ≤ 0.05) are colored red and increase and decrease in protein abundances are shown as ↑+M and ↓+M respectively. The specific numbers of protein expression changes from the analysis of *Fusarium DS 682*’s proteome in each treatment group is shown in Fig. 3a.

**Fig. S12.**
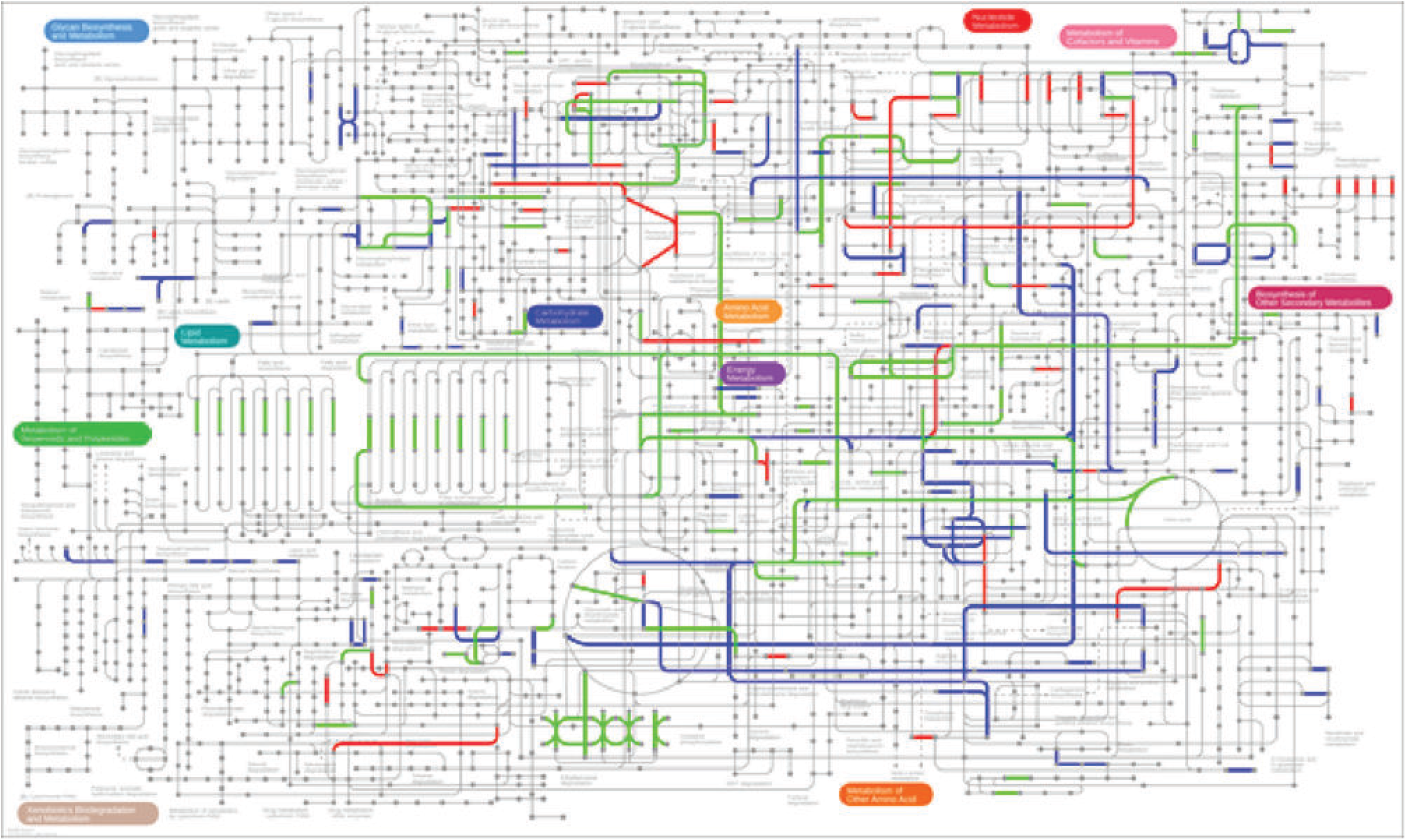
Several metabolic transport pathways are shifted under +M and -M growth conditions. Pathways highlighted in red are unique to +M, blue are unique to -M, and green are proteins associated with shared active KEGG metabolic pathways. Using KEGG enrichment analysis, we show that ATPases specific to oxidative phosphorylation pathways are measurable in fungal growth in both the presence and absence of minerals. However, there are specific ATPases, as shown in Fig. 3c and 4, that are enriched in fungal growth only in the presence of minerals that are indicative of increased transport of mineral nutrients. A more resolved KEGG map, that is navigable, can be found in iPath_3_ map here: <u>Fusarium sp. DS 682 Fungal Mineral Metabolic Pathway Module Transport</u>.

